# Establishment of Multi-stage Intravenous Self-administration Paradigms in Mice

**DOI:** 10.1101/2020.11.25.398503

**Authors:** Lauren M. Slosky, Andrea Pires, Yushi Bai, Nicholas Clark, Elizabeth R. Hauser, Joshua D. Gross, Fiona Porkka, Yang Zhou, Xiaoxiao Chen, Vladimir M. Pogorelov, Krisztian Toth, William C. Wetsel, Lawrence S. Barak, Marc G. Caron

## Abstract

A genetically tractable animal model would provide a needed strategy to resolve the biological basis of drug addiction. Intravenous self-administration (IVSA) is the gold standard for modeling cocaine and opioid addiction in animals, but technical limitations have precluded the widespread use of IVSA in mice. Here, we describe the first IVSA paradigms for mice that capture the multi-stage nature of the disease and permit predictive modeling. Mice with long-standing indwelling jugular catheters engaged in cocaine or opioid-associated lever responding that was fixed ratio- and dose-dependent, extinguished by the withholding of drug, and reinstated by the presentation of paired cues. Machine learning revealed that vulnerability to drug seeking and relapse were predicted by a mouse’s *a priori* response to novelty, sensitivity to drug-induced locomotion, and drug-taking behavior. Application of this behavioral and analysis approach to genetically-engineered mice will facilitate the identification of the neural circuits driving addiction susceptibility and relapse and focused therapeutic development.

## Introduction

Drug overdose is the leading cause of injury-related mortality in the U.S., claiming more than 70,000 lives in 2017 alone (*1*) and costing an estimated $78.5 billion per year in healthcare, lost productivity, and criminal justice involvement (*2*). While synthetic opioids, including fentanyl and its derivatives, are responsible for the spike in overdose deaths in recent years, mortality associated with cocaine and methamphetamine use disorders also tripled between 2012 and 2016 (*3*). Available therapies for opioid use disorders have limited efficacy, and there are no Food and Drug Administration (FDA) - approved therapies for the treatment of stimulant use disorders. This paucity of treatments persists despite a medical consensus that opioid and stimulant addictions are brain disorders and, as such, should be amenable to pharmacological interventions (*4*). The lack of effective therapeutics may be due to chemical addictions being a family of disorders whose multiple etiologies remain ill-defined. An understanding of the genetic and molecular determinants of addiction-associated behaviors will facilitate the identification of therapeutic targets and at-risk individuals.

Animal studies permit levels of control and manipulations that are not feasible with human subjects. Multiple paradigms, with varying levels of complexity, exist for modeling the effects of drug use in animals. Intravenous self-administration (IVSA) with operant conditioning is the gold standard for studying the effects of stimulants (e.g., cocaine, methamphetamine) and opioids (e.g., morphine, heroin, fentanyl) in animals. Pioneered in non-human primate and rat studies (*5-9*), this approach has proven to be the most translationally relevant addiction model, having greater face validity than locomotor and place preference protocols and demonstrated predictive validity (*10-12*).

The vast majority of IVSA studies conducted over the last 50 years have continued to use non-human primates and rats. Far fewer studies have utilized mice. Compared to other mammalian species, the mouse genome is the most amenable to genetic engineering. As such, many genetic tools, including global, conditional, inducible, and conditional-inducible transgenic strategies are available in mice. Other techniques that are well-developed in mice, but not other mammals, include optogenetic and chemogenetic approaches; strategies that permit the direct, real-time assessment of specific cell types and neural circuits to behavior.

The relatively fewer number of mouse IVSA studies to date are due, in part, to the technical challenges of applying this model to mice (*13*). Advances in materials, surgical techniques, and operant protocols, however, have led recently to unprecedented success in mouse self-administration approaches. While studies evaluating genetically-modified mice are becoming more common (*14-20*), IVSA in mice continues to be limited by low rates of study completion (e.g., 20-30% of trained mice), limited data collection (e.g., evaluation of a single drug dose on a single delivery schedule), and abbreviated study length (*16, 21, 22*).

Few studies to date have employed a longitudinal design in which acquisition, maintenance, extinction, and reinstatement of self-administration are assessed within the same cohort of mice. Longitudinal, multi-stage studies afford several advantages over past approaches, including the opportunity to describe the relationship between risk factors (e.g., trait constructs, behavioral signatures in *a priori* behavioral testing) and the development of addiction-associated behaviors (e.g., drug taking, drug seeking). Moreover, because of the chronicity of addictive behaviors, with discrete stages associated with distinct neurobiological determinants (*23*), longitudinal studies better approximate the clinical disease trajectory.

The primary objectives of the present study were two-fold. One goal was to develop multi-stage intravenous (iv) stimulant and opioid self-administration paradigms in mice. A second purpose was to use statistics to evaluate the relationships among observed variables and advance our understanding of their underlying behavioral constructs. To achieve these objectives, we evaluated self-administration of the psychostimulant cocaine and the short-acting opioid remifentanil. Here, we demonstrate that iv cocaine or remifentanil self-administration acquisition, maintenance, extinction, and reinstatement can be studied longitudinally in large cohorts of mice and provide a detailed description of drug-reinforced behavior in these paradigms. Exploratory factor analysis revealed that individual mouse performance during both the cocaine and remifentanil self-administration paradigms was a function of two latent variables that were differentially associated with *a priori* novelty-and drug-induced behavior in the open field. Application of machine learning approaches allowed us to develop predictive models of drug seeking and relapse. Critically, our approach to the behavioral and statistical analyses can be readily applied to genetically-engineered mice and adapted for the study of other drugs of abuse under diverse operant protocols.

## RESULTS

### Jugular catheterization and maintenance of catheter patency

IVSA requires the placement and maintenance of indwelling jugular catheters. In our optimized paradigm, post-surgical survival was high (i.e., 84% on post-surgery day 7; **Fig 1A**) and jugular catheter placement was highly effective (i.e., 98% patent on post-surgery day 7; **Fig 1B**). Throughout the duration of the study, attrition due to poor health was modest, with more than 70% of animals surviving to study completion or to catheter patency loss (**Fig 1C**). The catheters proved to be long-lasting, with a patency half-life of ≥ 100 days (**Fig 1D**). These data suggest that, with optimized materials and handling, jugular catheters can be placed reliably and maintained over long periods in mice. The newly established longevity of the animals and the catheters permitted us to adopt a longitudinal design for the study of iv cocaine and remifentanil self-administration acquisition, maintenance intake, extinction, and cue-induced reinstatement in the same mice.

**Figure 1.**
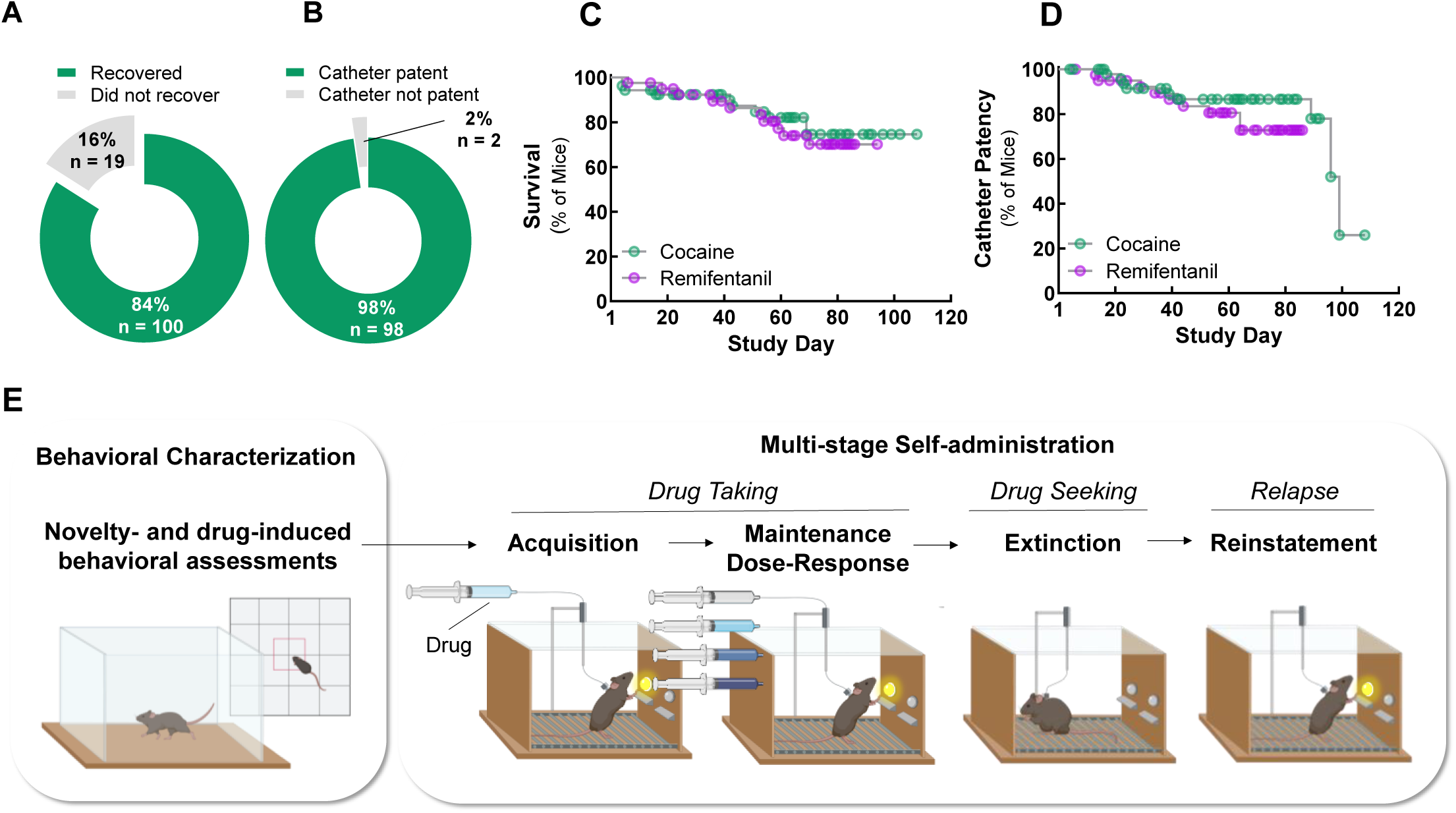
Long-lasting jugular catheters and low attrition permit longitudinal, multi-stage intravenous self-administration in mice. (A) Recovery from jugular catheterization procedure. Percent and number (n) of all animals that met survival and health criteria on post-catheterization day 7. (B) Success of jugular catheter placement. Percent and number (n) of all animals that passed a test of catheter patency on post-catheterization day 7. (C) Attrition over the study due to animals meeting designated health endpoints. Animals were removed from the study when they displayed signs of deteriorating health. The median time in the study (i.e., survival) exceeded 100 days. (D) Attrition over the study due to loss of catheter patency. Cather patency was followed throughout the study. The catheters of animals removed from the study for health reasons were considered patent unless they had a documented patency test failure. The median catheter patency life was ≥ 100 days. (E) Overview of study design. Mice with indwelling jugular catheters underwent behavioral testing in the open field and were then trained to self-administer drugs paired with a cue light via lever responding in operant chambers. For the details of the contingent advancement training and testing paradigms, see Figures S1 and S2.

### Acquisition of cocaine and remifentanil self-administration

Following catheter placement and prior to self-administration training, novelty-induced exploratory behaviors and cocaine (20 mg/kg, i.p.)-or remifentanil (0.1, 1 and 10 mg/kg, i.p.) - induced locomotor activities were assessed in the open field (**Fig 1E, S3**). Like cocaine (**Fig S3A**) and other opioids, remifentanil produced transient dose-dependent increases in locomotor activity, consistent with its short half-life (**Fig S3B**). Mean values were within the anticipated ranges for all parameters.

Mice with patent indwelling jugular catheters were trained to self-administer cocaine (0.5 mg/kg/infusion, iv) or remifentanil (0.1 mg/kg/infusion, iv) paired with a cue light by lever responding in operant chambers. We established contingent advancement protocols (**Fig S1, S2**) to address the challenges of behavioral inconsistency and high inter-mouse variability. Our protocols used predominantly short access, 1-hour daily sessions to reduce the required chamber time per mouse per day and, thereby increasing the number of animals that could be tested as a cohort.

Eighty-four percent and 94% of mice with patent catheters acquired cocaine (0.5 mg/kg/infusion) and remifentanil (0.1 mg/kg/infusion) self-administration, respectively (**Fig 2A**,**C**); this percent did not differ by drug class (**Table S1**). The mean session number required to meet final acquisition criteria was lower for remifentanil (10.8 ± 0.2 sessions) than for cocaine (13.5 ± 0.6 sessions), although the peaks of frequency histogram curves were indistinguishable (**Fig 2B**,**D, Table S1**). Notably, variability in this time-to-train measure was significantly lower for remifentanil relative to cocaine (F(36, 32) = 9.077, p<0.001, F test for variances), suggesting that, at these doses, remifentanil may be the more effective reinforcer.

**Figure 2.**
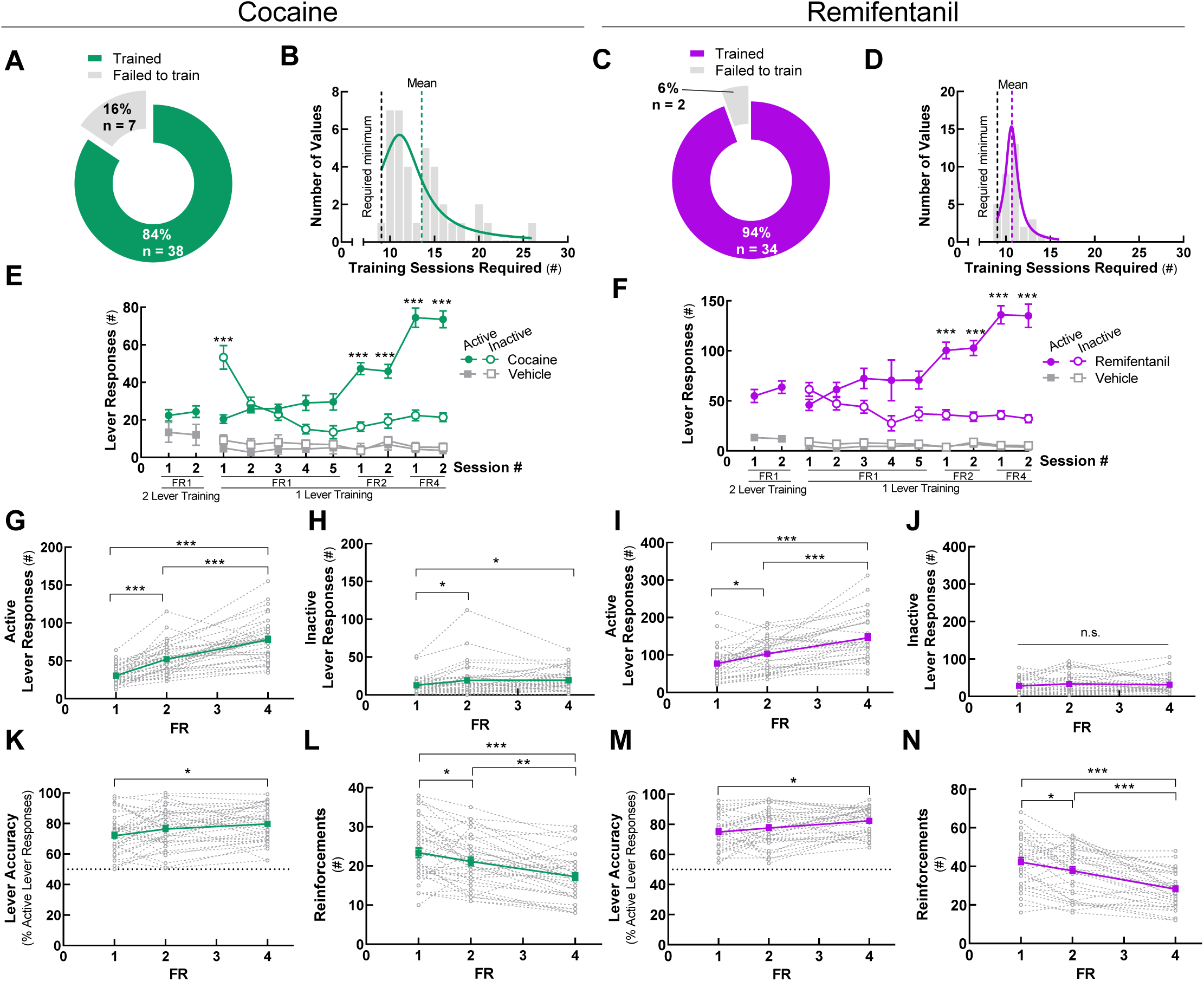
Acquisition of cocaine and remifentanil self-administration. Mice with indwelling jugular catheters were trained to self-administer iv cocaine (0.5 mg/kg/infusion) or remifentanil (0.1 mg/kg/infusion) via lever responding in operant chambers at FR1, FR2, and FR4. Group means are represented by *solid green* (cocaine) or *purple* (remifentanil) *lines* and individual replicates by *dashed lines*. Note, *solid gray lines* represent the vehicle. (A, C) Acquisition success rates. Percent of animals with patent catheters that acquired or failed to acquire iv cocaine (A) or remifentanil (C) self-administration at FR4 using contingent advancement protocols. (B, D) Sessions to self-administration acquisition. Frequency plot of the number of sessions required to meet final cocaine (B) or remifentanil (D) self-administration acquisition criteria at FR4. The minimum session requirements and group means are indicated by *dashed vertical lines*. (E, F). Lever responses by acquisition session type. Lever responses by mice that acquired cocaine (E) or remifentanil (F) self-administration. Data are presented from the first sessions of each session type, as indicated. (G, I) Active lever responses. Total active lever presses by mice self-administering cocaine (G) or remifentanil (I) at FR1, FR2, and FR4. Responses include both those that contributed to earned reinforcements and those that occurred during the post-reinforcement time-out period. (H, J) Inactive lever responses. Total inactive lever presses by mice self-administering cocaine (H) or remifentanil (J) at FR1, FR2, and FR4. (K, M) Lever accuracy. Percent of total lever responses that occurred at the designated active lever by mice self-administering cocaine (K) or remifentanil (M) at FR1, FR2, and FR4. (L, N) Reinforcements. Earned reinforcements by mice self-administering cocaine (L) or remifentanil (N) at FR1, FR2, and FR4. For details on statistical comparisons, see **Tables S1, S2**. Also see **Figures S1, S2**.

Because cues themselves can be reinforcing (*24*) and some mice initially display high rates of responding in the absence of a reward (*19, 20*), specificity of lever responding for the drug reinforcers was assessed. Like mice in the cocaine and remifentanil reinforcement groups, animals in a vehicle reinforcement group initially engaged in moderate levels of lever responding. However, they failed to develop active *versus* inactive lever discrimination, and their lever responding was unaffected by increasing the FR schedule (**Fig 2E**,**F**). Together, these data demonstrate that iv cocaine and remifentanil self-administration can be readily and rapidly acquired in short, daily sessions by mice without prior operant training.

### Cocaine and remifentanil self-administration are fixed-ratio-dependent

As the response requirement to earn a reinforcement increases, the extent of self-administration decreases. This inverse relationship generalizes across species and drug classes (*25*). As described for rats, non-human primates, and humans, self-administration behaviors in mice changed as the FR schedule was increased. In both drug paradigms, active lever responses increased from FR1 to FR2 and from FR2 to FR4 (**Fig 2G**,**I**). During these transitions, inactive lever responding increased only modestly for cocaine (**Fig 2H**) and remained unchanged for remifentanil (**Fig 2J**). Consequently, lever discrimination (i.e., accuracy) increased between FR1 and FR4 (**Fig 2K**,**M**). At FR4, 80 ± 2% and 82 ± 2% of lever responses occurred at the active, drug-delivering lever for cocaine and remifentanil, respectively. The increase in active lever responding with the increasing FR schedule was not sufficient to maintain levels of drug intake. The number of self-administered reinforcements of both drugs was reduced at FR4 compared to FR1 and FR2 (**Fig 2L**,**N**). These results demonstrate that the number and percent of active lever responses, and therefore drug intake, is a function of the FR schedule in mice. The number of reinforcements earned per hour and the amount of cocaine or remifentanil intake at FR1 is consistent with published results for mice trained using other paradigms, which have used different acquisition protocols and longer sessions (*16*).

### Cocaine and remifentanil self-administration are dose-dependent

Once cocaine or remifentanil self-administration was acquired at FR4 at the training doses, we systematically varied the dose delivered per infusion. We assessed stable lever responding at 5 cocaine doses (0.1, 0.3, 0.5, 1.0, and 3.0 mg/kg/infusion) and 6 remifentanil doses (0.01, 0.03, 0.1, 0.3, 1.0, and 3.0 mg/kg/infusion), permitting of the establishment of dose-effect curves. As demonstrated previously in rats and non-human primates (*9, 26, 27*), there was a clear drug dose-dependency to self-administration responses. For both cocaine and remifentanil, there was a characteristic inverted U-shape relationship (*25, 28*) between the earned reinforcements relative to the log drug dose (**Fig 3A**,**I**) and active lever responses relative to log drug dose (**Fig 3B**,**J**). Earned reinforcements did not differ by sex (**Fig S4**). The cocaine intake vs. log cocaine dose data were fit to a sigmoidal curve (**Fig 3D**) and the remifentanil intake vs log remifentanil dose data were fit to an exponential growth curve (**Fig 3L**). Notably, inactive lever responding for both drugs was also drug dose-dependent, decreasing linearly with the log cocaine or remifentanil dose (**Fig 3C**,**K**). Lever accuracy increased linearly with log drug dose for both drugs (**Fig 3E**,**M**). Two latency measures, latency to initiate lever responding and latency to earn the first reinforcement, are considered indices of urgency or motivation to initiate drug taking (*29, 30*). While the latency to the first lever response within a session was unaffected by drug dose (**Fig 3F**,**N**), the latency to the first reward decreased linearly with log drug dose (**Fig 3G**,**O**). The number of lever responses that occurred during the post-reinforcement time-out periods, a parameter that may be related to action impulsivity (*31*), was also drug dose-dependent, as it decreased linearly with the log cocaine and remifentanil dose (**Fig 3H**,**P**). This time-out responding vs. log drug dose relationship remained significant after normalizing the data to total lever responding, and thus the length of the time-out interval, for cocaine but not for remifentanil (**Fig S5**). Total earned reinforcements and active lever responses were higher and the latency to initiate drug taking was lower in the remifentanil paradigm, suggesting that, in these dose ranges, remifentanil may be a stronger reinforcer than cocaine.

**Figure 3.**
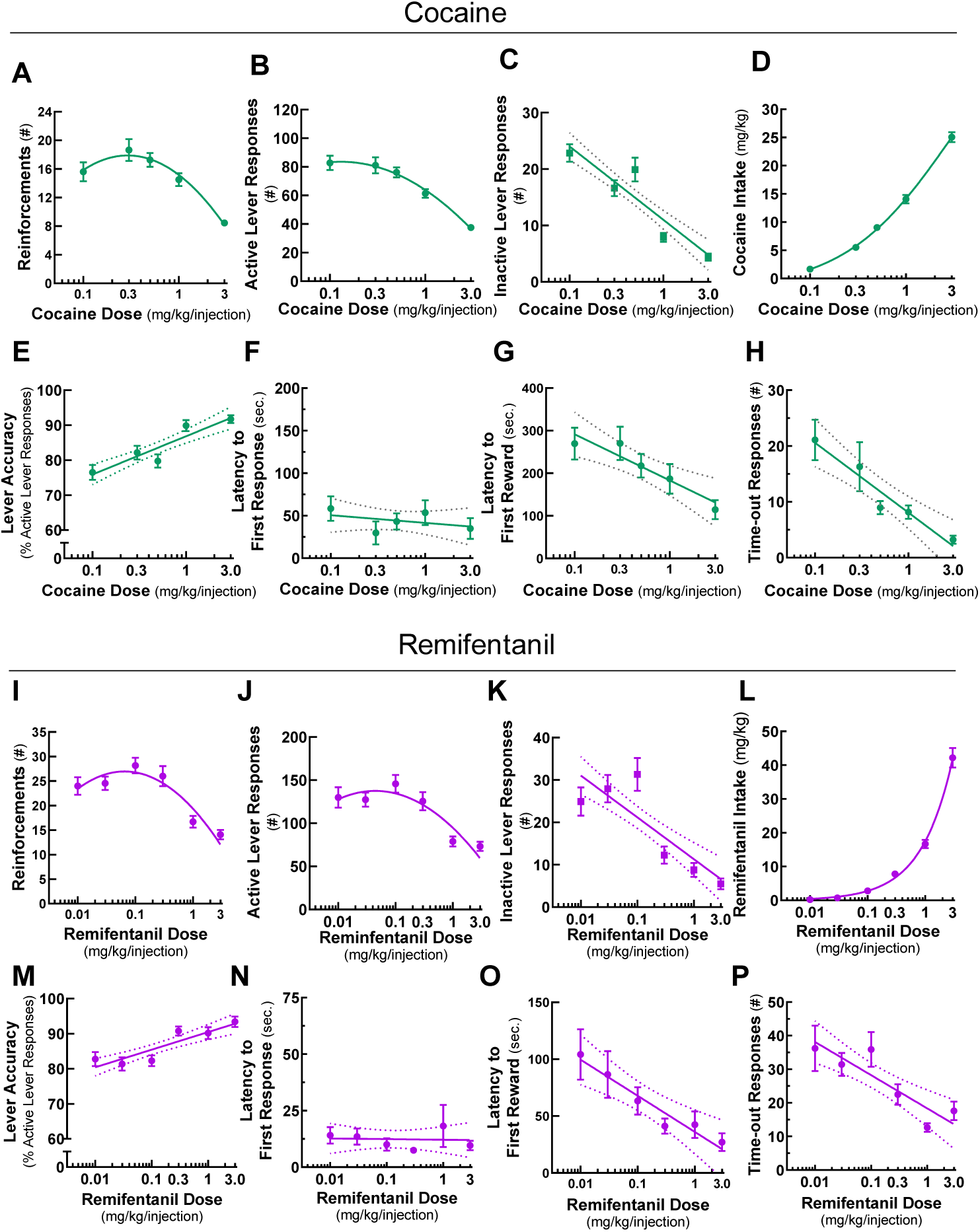
Dose-response relationships of cocaine and remifentanil self-administration behaviors. Stable self-administration of cocaine at FR4 was assessed in 60 min sessions at doses of 0.1, 0.3, 0.5, 1.0, and 3.0 mg/kg/infusion. Stable self-administration of remifentanil at FR4 was assessed in 60 min sessions at doses of 0.01, 0.03, 0.1, 0.3, 1.0, and 3.0 mg/kg/infusion. Ranges of axes were selected to emphasize curve fits and differences by drug class. (A, I) Reinforcements. Earned reinforcements vs. log cocaine (A) or remifentanil (I) dose. Data were fit to a second order polynomial curve. (B, J) Active lever responses. Active lever presses vs. log cocaine (B) or remifentanil (J) dose. Responses include both those that contributed to earned reinforcements and those that occurred during the post-reinforcement time-out period. Data were fit to a second order polynomial curve. (C, K) Inactive lever responses. Inactive lever presses vs. log cocaine (C) or remifentanil (K) dose. Data were fit to a line. (D, L) Drug intake. Cocaine (D) or remifentanil (L) consumed vs. log drug dose. Cocaine data were fit to a sigmoidal curve. Remifentanil data were fit to an exponential growth curve. (E, M) Lever accuracy. Percent of total lever responses that occurred at the designated active lever vs. log cocaine (E) or remifentanil (M) dose. Data were fit to a line. (F, N) Latency to first lever response. Time from the start of the session until the first lever response vs. log cocaine (F) or remifentanil (N) dose. Data were fit to a line. (G, O) Latency to first earned reinforcement. Time from the start of the session until the first earned reinforcement vs. log cocaine (G) or remifentanil (O) dose. Data were fit to a line. (H, P) Post-reinforcement time-out responses. Total number of lever responses that occurred during the post-reinforcement time-out periods vs. log cocaine (H) or remifentanil (P) dose. Data were fit to a line. For information on curve fits, see **Table S3**. Also see **Figures S4, S5, S6**

To investigate the kinetics of self-administration during the 1-hour sessions, we constructed cumulative reinforcement time courses for all drug doses (**Fig S6**). These plots were linear for all remifentanil doses and for cocaine up to the 1.0 mg/kg/infusion dose, indicating that self-administration occurred at a constant rate over the course of the sessions. Together, these results demonstrate that mice exhibit dose-dependent self-administration behaviors.

### Cocaine-and remifentanil-associated lever responding can be extinguished and reinstated

It has been demonstrated in rats and non-human primates (*32-34*), and, more recently, in mice (*17, 24, 35, 36*) that drug-associated lever responding can be extinguished over repeated sessions where no drug is available, drug-associated cues are withheld, and lever responses have no programmed consequences - a process known as extinction (*37*). After mice completed dose-response testing, they were subjected to once daily, 1-hour extinction sessions (**Fig S1, S2**). In the cocaine paradigm mice were extinguished to a criterion (i.e., ≤ 20% of active lever responses for a 1.0 mg/kg/infusion of cocaine), and in the remifentanil paradigm all mice completed 20 extinction sessions regardless of performance. For both drugs, active lever responding decreased as a function of extinction session number (**Fig 4A**,**E**), with mice requiring 20 ± 1.6 sessions to meet extinction criteria for cocaine. The reduction in total lever responding over the first 20 extinction sessions can be described using a one-phase exponential decay function (**Fig 4B**,**F**). Notably, the rate of extinction was comparable for both drugs. From the final 1.0 mg/kg/infusion self-administration session to the first extinction session, lever responding was increased (**Fig 4A**,**E**) and lever accuracy was reduced (**Fig 4C**,**G left**), suggesting the mice actively seek drug in this session. By the final extinction session, lever discrimination was lost in the cocaine group (**Fig 4C, middle**) and was diminished in the remifentanil group (**Fig 4G, middle**).

**Figure 4.**
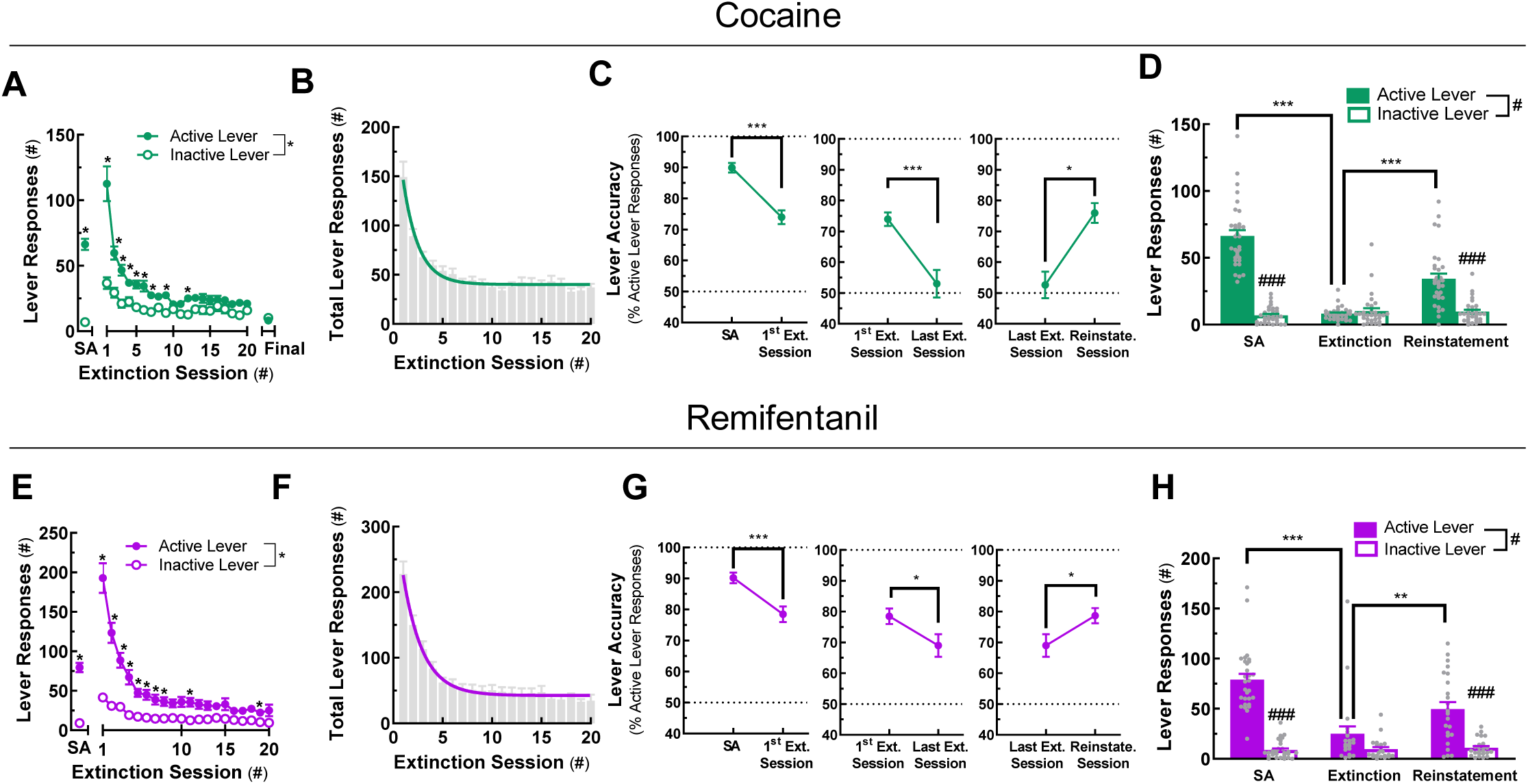
Extinction and cue-induced reinstatement of cocaine- and remifentanil-associated lever responding. In mice trained to self-administer cocaine or remifentanil at FR4, drug-associated lever responding was extinguished in consecutive sessions in which drugs and cues were withheld and subsequently reinstated by the reintroduction of drug-paired cues. (A, E) Active and inactive lever responses during extinction. Active and inactive lever responding is presented by extinction session number for mice in the cocaine (A) and remifentanil (E) paradigms. Lever responding for 1 mg/kg/infusion of drug (i.e., self-administration, SA) is included for reference. (B, F) Total lever responses during extinction. Total lever responding is presented by extinction session number in *gray bars* for mice in the cocaine (B) and remifentanil (F) paradigms. *Colored curves* are one-phase exponential decay fits. (C, G) Lever accuracy at maintenance SA, at the final extinction session, and at reinstatement. The percent of total lever responses occurring at the active lever is presented by session type for mice in the cocaine (C) and remifentanil (G) paradigms. SA indicates accuracy during the 1.0 mg/kg/infusion drug maintenance active self-administration session. (D, H) Active and inactive lever responses during reinstatement. Active and inactive lever responding is presented by session type for mice in the cocaine (D) and remifentanil (H) paradigms. SA indicates lever responding during the 1.0 mg/kg/infusion drug active self-administration session. Group means are represented by *colored bars* and individual values are shown as *gray circles*. For curve parameters and details on statistical comparisons, see **Table S4**.

Presenting drug-experienced animals or humans with paired contextual or environmental cues can reinstate drug seeking (*38*), a phenomenon known clinically as relapse. In mice that completed extinction, re-introduction of drug-associated cues, including the cue light, syringe pump sounds, and lever movements, increased responding at the previously active lever and reinstated lever discrimination (**Fig 4C, right, 4G, right, D**,**H**). These results suggest that mice trained to self-administer cocaine or remifentanil in once daily, 1-hour sessions display characteristic operant behaviors observed in other paradigms and species. Hence, cocaine- and remifentanil-associated lever responding in mice with restricted daily access to drug can be extinguished by withholding the drug and reinstated by presentation of the drug-associated cues.

### Dimension reduction of self-administration data identifies latent variables

Collecting multiple measures from a given subject and having groups of sufficient size permits the use of multivariate analysis. Our data provide an ideal large data set (i.e., n > 30 mice per condition) to assess the latent variables underlying drug taking and seeking in mice.

Self-administration behaviors both within and across sessions were related. These relationships were visualized in correlation matrix heat maps and dendrograms for 11 key variables from the cocaine and remifentanil paradigms (**Fig 5, Table S5, S6**). These variables included measures of drug taking averaged across the dose-response curve (i.e., self-administered reinforcements, active and inactive lever responding, lever accuracy, and latency measures), as well as indices of drug seeking (i.e., lever responding during extinction and cue-induced reinstatement). The internal structures of the drug taking-associated behavioral measures were remarkably similar in the cocaine and remifentanil paradigms (**Fig 5**, variables 1-8). Differences between drug classes were apparent, however, in the relationships between the drug seeking measures (**Fig 5**, variables 9-11).

**Figure 5.**
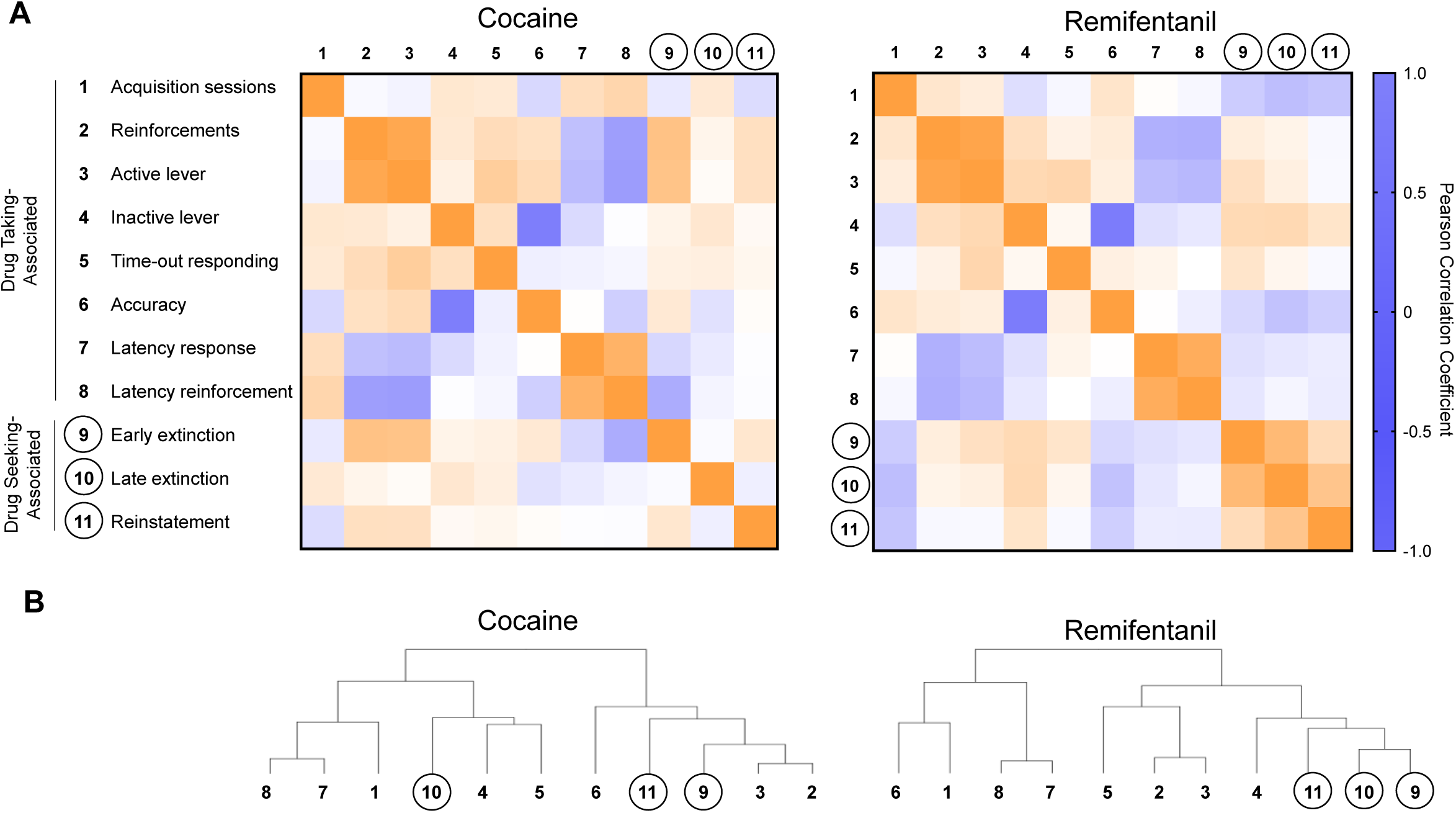
Relationships between drug taking- and drug seeking-associated self-administration variables. Drug taking- and drug seeking-associated variables are numbered according to the key on the far left. Numbers assigned to drug seeking-associated variables are enclosed in circles. (A)Heat maps of Pearson R correlation coefficients and corresponding p-values. The increasing saturation of *orange* and *blue color* correspond to increasing absolute values of positive and negative Pearson correlation coefficients, respectively. The increasing saturation of green color corresponds to decreasing p-values. *White* indicates a correlation coefficient close to 0 and p-value close to 0.4. (B) Dendrograms. Results of hierarchical clustering. Correlation coefficient values and corresponding p-values are provided in **Tables S5** and **S6**.

For interrelated datasets such as these, dimension reduction strategies are useful. Such strategies may identify a reduced number of latent, orthogonal constructs contributing to common variance. Indeed, this approach has previously been used to reduce the dimensionality of open field data in mice (*39*) and of self-administration and locomotor activity data in rats (*40*). Bartlett’s Test of Sphericity indicated that there was sufficient shared variance between the selected measures in **Fig 5** to warrant dimension reduction.

We conducted two within-drug exploratory factor analyses (EFAs) with 12 and 11 variables from the cocaine and remifentanil datasets, respectively (**Fig 6, Table S7**). We extracted 2 factors explaining a cumulative 52% of the total variance of the model for cocaine and 55% of the total variance of the model for remifentanil (**Fig 6, Table S7**). In loading plots (**Fig 6A**,**B**), the more variance within a measure that is explained by a given factor, the higher the factor score and the further from the origin the variable is located. Variables that are correlated will load onto a common factor, with variables that are directly proportional loading onto the same side of the origin, and variables that are inversely proportional loading onto opposite sides of the origin.

**Figure 6.**
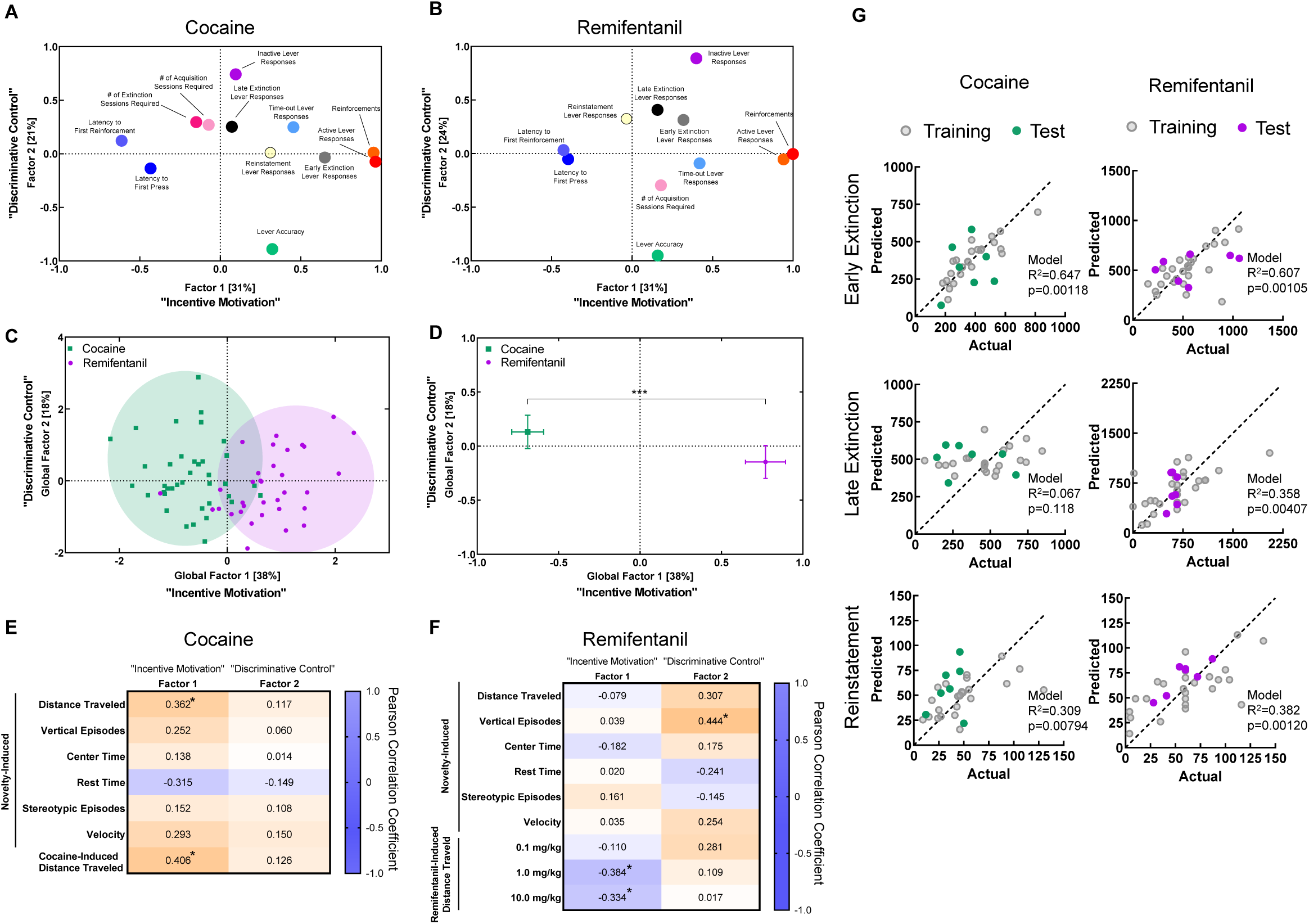
Feature discovery and predictive modeling of self-administration behaviors. (A,B) Variable factor loading. Factor loading plots for self-administration variables included in exploratory factor analyses for cocaine (A) and remifentanil (B). (C) Global factor scores. Factor scores for individual animals, obtained using the regression method, are represented as *squares* or *circles. Ovals* are drawn around animals in each drug/reinforcer paradigm. Each oval encompasses the genotype mean and contacts the closest 80% of individual values. (D) Comparison of factor scores by reinforcer. Factor scores for components 1 and 2 were averaged by drug paradigm and compared using unpaired, two-tailed student’s t-tests. For component 1: t(70)=9.4, p<0.0001. For component 1: t(70)=1.3, p=0.2082. (E,F) Heat maps of Pearson correlation coefficients for open field variables and factor scores. Correlation coefficients for cocaine (E) and remifentanil (F) are shown on top for heat maps in which the increasing saturation of *orange* and *blue color* correspond to increasing absolute values of positive and negative Pearson correlation coefficients, respectively. Significant values are indicated by an asterisk. *p<0.05. (G) Actual vs. predicted plots for multiple linear regression models constructed for drug seeking during early extinction (top), late extinction (middle), and reinstatement (bottom) based on responses to novelty, drug-induced hyperlocomotion, and drug-taking behaviors. For factor loadings, see **Table S7**. For information on variables included in the regression models and details on model performance, see **Tables S8, S9** and **S10**, and **Figure S8**.

The first factor extracted accounted for 31% (eigen value = 3.728) of the cocaine model and 31% (eigen value = 3.396) of the remifentanil model. Based on variable loading, we determined that this factor represented an incentive motivation construct. The same variables (i.e., active lever responses, earned reinforcements, latencies to first lever response, and first reinforcement during self-administration) loaded most heavily onto this factor in both models (**Fig 6A, Table S7**). The positions of these loadings indicate that animals with higher active lever responses and earned reinforcements had lower latencies to their first lever response and reinforcement.

The second factor extracted accounted for 21% (eigen value = 2.562) of variance in the cocaine model and 24% (eigen value = 2.673) of variance in the remifentanil model. We determined that factor 2 represented a discriminative control dimension, as the variables that loaded most heavily on this factor in both models were lever accuracy and inactive lever responding during self-administration (**Fig 6A, Table S7**). As expected, this finding indicates that animals with lower lever accuracy produced more responses at the inactive lever during self-administration.

We next examined how drug seeking during extinction and cue-induced reinstatement loaded onto these factors. For cocaine, while early extinction lever responding was well-explained by the incentive motivation factor 1, late extinction responding was not. Instead, late extinction responding loaded weakly on the discriminative control factor 2. For remifentanil, early and late drug seeking clustered together, but were only modestly well-explained by either factor. Interestingly, cue-induced reinstatement of cocaine seeking loaded exclusively on factor 1, while that of remifentanil seeking loaded exclusively on factor 2. Together, the correlation matrices and factor analysis suggest that while patterns of cocaine and remifentanil *taking* are similar those of cocaine and remifentanil *seeking* are distinct. Drug seeking during early extinction, late extinction, and cue-induced reinstatement may be driven by distinct underlying biological and/or behavioral programs that differ by drug class.

### Identification of drug class-specific self-administration phenotypes

To determine whether cocaine and remifentanil produced distinct self-administration phenotypes, a global exploratory factor analysis was performed with mice from both drug paradigms (**Fig S7, Table S7**). Individual mice were assigned regression factor scores (**Fig 6C**) and factor scores were compared between reinforcers. Animals in the cocaine and remifentanil paradigms differed on incentive motivation factor 1, but not on factor 2 (**Fig 6D**). This finding demonstrates that EFA can identify drug class-specific self-administration phenotypes.

### Predicting self-administration performance from *a priori* open field assessments

Because self-administration is time and resource-intensive, it would be desirable to identify predictors of self-administration behaviors in other, more facile behavioral assays, including novelty- and drug-induced motor activities in the open field. As such, we assessed whether there was a relationship between the *a priori* exploratory or drug-induced behaviors in the open field and self-administration performance, as determined by factor 1 and 2 scores (**Fig 6E**,**F**). Drug-induced locomotion was positively and negatively associated with factor 1 for cocaine and remifentanil, respectively. Exploratory vertical episodes (i.e., rearing) correlated with factor 2 scores for remifentanil, but not for cocaine. These findings suggest that the relationship between open field motor activities and self-administration factor scores differ by drug class.

### Predicting drug seeking from *a priori* open field assessments and drug taking behavior

Drug seeking during extinction and reinstatement are the most clinically-relevant paradigm stages, but they were not well described by the factors extracted in the EFA. To examine whether drug seeking during early extinction, late extinction, or reinstatement can be predicted by a mouse’s response to novelty, drug-induced hyperactivity, or self-administered drug intake, we applied machine learning algorithms. To remove multicollinearity, we performed principal component analyses on novelty-induced open field (**Table S8**) and drug-taking behaviors (**Table S9**). We then divided the data into training and test datasets using a random 80:20 split and built mulitple regression models based on the novelty-response component scores, drug-taking component scores, and drug-induced hyperlocomotion data of the training set. Predictive models were successfully generated for both drugs and all drug seeking stages, except for late extinction responding for cocaine (**Figure 6G, Table S10**). Residuals for the training and test sets were comparable for the late extinction and reinstatement models and only modestly higher for the early extinction models (**Figure S8**), suggesting these models are not overfit. Interestingly, the variables contributing predictive value to the models differed both by drug and drug-seeking stage. These data indicate that drug seeking can be predicted from a mouse’s response to novelty, sensitivity to drug-induced locomotion, and drug taking behavior, but that the relative importance of these measures differs by drug type and drug-seeking stage.

## DISCUSSION

Causal studies on the biological basis of drug addiction are not practical or ethically achievable in humans. Thus, understanding disease mechanisms requires animal models. The most versatile and practical models have been rats. Technical obstacles have largely precluded longitudinal, multi-stage self-administration studies in mice, the mammalian species for which the most genetic tools are available. Here, we have overcome historic obstacles and developed expeditious paradigms for cocaine and remifentanil in which mice with indwelling jugular catheters progress through the self-administration stages of acquisition, maintenance, extinction, and reinstatement in a systematic manner that permits comparison of behavioral parameters across drug class and the application of predictive modeling.

IVSA requires placement of catheters into the jugular vein. Historically, jugular catheter placement, maintenance of catheter patency, and prevention of infection secondary to their use have limited the extent and duration of self-administration studies in mice (*13*). The jugular catheterization procedure in mice is not trivial, due in part to their small size (*13*). We used a guide needle technique modified from (*41*), and used polyurethane catheters that are less thrombogenic than those composed of polyethylene or polyvinyl chloride. The new-found longevity of surgically-implanted mice and catheters in the current paradigms has permitted longitudinal, multi-stage study designs.

In many self-administration paradigms, drug exposure is preceded by operant conditioning with food (*24, 42*). To reduce the total time required for these types of studies, circumvent potential confounding effects of prior conditioning (*24*), and permit the future assessment of genotype differences in self-administration acquisition, we did not conduct prior operant training. In acquisition sessions, mice progressively learned active vs. inactive lever discrimination and self-administered drugs paired with a cue light on FR1, FR2, and FR4 schedules. The extent of self-administration during this training period was consistent with publications for the dose, time frame and reinforcement requirements in mice for cocaine (*16, 41*) and remifentanil (*43*). The observed increase in total response output with increased response requirement is a hallmark of drug self-administration across species and drug class (*25*), and it was not observed in mice given access to the vehicle rather than cocaine or remifentanil. The percent of mice achieving final cocaine self-administration acquisition criteria (∼80%) was consistent with that previously reported for C57BL/6J mice, and it is significantly higher than that reported for other mouse strains (*16, 21*). In our experiments, mice achieved high levels of lever discrimination during training for both drugs, as evidenced by the percent of total responses occurring at the active lever.

Once cocaine or remifentanil self-administration was achieved, animals progressed through a maintenance dose-response phase in which the drug dose per infusion was systematically varied. For both cocaine and remifentanil, the number of reinforcements earned per hour and the extent of drug consumption across doses was consistent with published results (*16, 41, 43*). Interestingly, except for latency to the first response, all behavioral indices showed dose-dependence for both drugs.

Following maintenance dose-response testing, animals moved into an extinction phase. Extinction is more protracted in mice than in rats and nonhuman primates (*24*). The rate and extent of extinction in the present study is consistent with previous reports for C57BL/6 and A/J mice extinguished in the absence of cocaine-associated cues (*24*). However, the extinction in the current study was more effective, in that mice achieved greater reductions in active lever responding than that reported in cocaine-conditioned C57BL/6J mice when extinction occurred in the presence of paired cues (*24*). This indicates that the presence of paired cues may hinder behavioral extinction.

As documented in multiple species, including mice (*17, 24*), lever responding in our paradigm decreased with successive extinction sessions in a manner that was best fit by a two-phase model: rapidly at first (i.e., sessions 1-4) and then more slowly (i.e., sessions 5-20). At the time mice met final extinction criterion, active lever responding was not only reduced, but active vs. inactive lever discrimination was lost for cocaine and significantly reduced for remifentanil. For cocaine, these results suggest an absence of goal-directed drug seeking and successful extinction of drug-associated lever responding.

Following extinction, mice completed a reinstatement session in which drug-associated cues were presented. This reintroduction of drug-paired cues reinstated both responding at the previously active lever and active vs. inactive lever discrimination. The finding that limited access to cocaine or remifentanil afforded by once daily, 1-hour sessions is sufficient to produce this behavior fits with emerging evidence from the rat literature suggesting that exposure to large quantities of cocaine is not necessary for escalation of intake, incentive-sensitization, or drug or cue-induced reinstatement of drug seeking behavior (*44*). Many reinstatement paradigms have been described, including those using various environmental cues, drug priming, and stress. Reinstatement of drug seeking induced by different stimuli (e.g., cue-induced, drug-induced, stress-induced) occurs through distinct mechanisms (*45*). We elected to use drug-paired cues as the stimulus for reinstatement in our study because of clinical reports demonstrating a relationship between drug-associated cues and relapse (*46*). In the future, this paradigm can be adapted for the study of reinstatement induced by other stimuli.

The longevity of the jugular catheters and mice in our experiments permitted a longitudinal study design. One advantage of employing a longitudinal design with a larger sample size is the ability to examine the relationships between behaviors at different time-points (*47*). This approach may be especially useful for self-administration studies, where later time-points, such as at the extinction and reinstatement stages, may be more clinically-relevant and more time-intensive, costly, and technically challenging to assess as compared to the earlier behavioral acquisition and maintenance phases. We performed a series of EFAs to examine how the variables examined in this study are related. We extracted two components from this analysis and, based on variable loading that was consistent between the drug paradigms, termed the one with the highest explanative value “ incentive motivation” and the one with the next highest explanative value “ discriminative control.”

Several aspects of the resulting model are informative. The first is the high degree of similarity between the internal structure of the drug-taking measures for cocaine and remifentanil. The second is the contrast between the structures of cocaine and remifentanil seeking data and their differential relationships to active and inactive lever responding during self-administration.

In the cocaine EFA, variable loadings support the idea that there are two distinct phases of lever responding during the extinction phase of the longitudinal self-administration paradigm. While early extinction responding is highly correlated with the extent of self-administration and urgency to initiate drug taking, late extinction responding is more closely associated with variation in lever discrimination and inactive lever responding during self-administration. A two-phase extinction model is supported further by the shape of the total presses vs. extinction session number best-fit curve and by prior observations in support of the early- and late-stage extinction phases as biologically distinct. For example, loss of β-arrestin2 in the infralimbic prefrontal cortex affects extinction responding only in its early stages (i.e., sessions 1-4), and not in subsequent sessions (*17*). Another noteworthy observation is that the number of sessions required to reach final extinction criteria in the cocaine-administered mice was almost entirely unrelated to the extent of self-administration during the maintenance stage, suggesting that the rate of extinction learning is independent of the degree of cocaine consumption. Conversely, the rate of extinction was closely associated with the rate of self-administration acquisition, consistent with the notion that acquisition and extinction are both learning processes (*48*).

Additionally, it is significant that late extinction responding in the cocaine cohort and both early and late extinction responding in the remifentanil cohort were related more closely to inactive lever responding than to active lever responding during the maintenance dose-response phase. At least two interpretations of these findings are possible. One is that lever responding in later extinction sessions does not reflect active drug seeking, but instead is a perseverative response manifested from poor action impulse control. An alternative interpretation is that responding at the inactive lever during the maintenance dose-response phase is not solely a reflection of discriminative control, but it is also an indication of active drug seeking. Drug seeking during the cue-induced reinstatement session was explained moderately by Factor 1 (incentive motivation) in the cocaine paradigm and by Factor 2 (discriminative control) in the remifentanil paradigm, suggesting that patterns of drug-seeking behaviors may differ by drug class.

A correlation between novelty-induced locomotor activity and vulnerability to psychostimulant but not opioid self-administration has been reported in rats (*49, 50*). Here, we document a parallel phenomenon in mice. Exploratory (i.e., novelty-induced) locomotor activity was positively correlated with the extent of cocaine self-administration, captured by cocaine EFA Factor 1, but not with the extent of remifentanil self-administration, captured by remifentanil EFA Factor 1. Notably, drug-induced hyperlocomotion is a positive and negative predictor of Factor 1 scores for cocaine and remifentanil, respectively.

Identifying at-risk individuals prior to the development of a chemical addiction is a major clinical goal. Here, we develop the first predictive models of maladaptive drug seeking behaviors in mice. Our ability to build such models based on reactions to novelty, response to acute drug exposure, and patterns of drug intake suggests relationships among these behaviors can be leveraged to identify at-risk subjects. The observation that the best predictors differed by drug and by drug seeking stage (e.g., early extinction vs. late extinction vs. cue-induced reinstatement) supports the results of the cluster analysis and EFA and indicates that cocaine and remifentanil seeking are driven by underlying biological and/or behavioral programs that are distinct and stage-specific.

While sex effects on self-administration behaviors have been reported in rats, particularly in long-access paradigms (i.e., >6 hr) (*51*), and in C57BL/6J mice on progressive ratio schedules (*52*), no such effect on self-administration was identified in the current FR-based paradigms. Our finding agrees with a report for cocaine self-administration (*52*), and others for alcohol (*53*) and food reinforcement (*54*), that sex differences are evident for C57BL/6 mice only when reinforcements are delivered on more behaviorally demanding schedules.

While the present work represents a significant advance in the length and extent of IVSA studies in mice, attrition over the study duration and inter-animal variability have not been eliminated. Moreover, self-administration is not acquired by all mice. As such, for these types of longitudinal studies, one should expect to start with 40-45 mice for each longitudinal arm to finish with the 30+ cohort size required for sufficient statistical rigor.

Together, the present findings analyze cocaine- and remifentanil-reinforced behaviors in longitudinal self-administration paradigms in which cohorts of mice progressed through the stages of acquisition, maintenance, extinction, and reinstatement. The systematic progression of mice through these paradigms allows comparisons to be made across genetically-engineered mouse lines, reinforcing substances, and potential anti-addiction treatments. Incorporation of other drug access schedules (e.g., intermittent access) and reinforcement schedules (e.g., progressive ratio) will expand its utility. Critically, through the application of this multi-stage self-administration paradigm to the large and growing number of genetically-engineered mice, we can now conduct clinically-relevant, detailed studies on the contributions of genes to addiction. The knowledge gained from these studies may facilitate the identification of at-risk individuals and permit the development of mechanism-based therapeutics for a variety of substance use disorders.

## MATERIAL AND METHODS

### Animals

This study employed adult male and female β-arrestin2^flox/flox^ mice on a C57BL/6J background. These inbred mice were selected for study because they could also serve as controls in conditional β-arrestin2 knock-out studies, the results of which will be presented elsewhere. At the start of studies, animals were 12-24 weeks old and 19–30 g body weight. Prior to self-administration testing, mice were group housed and maintained on a 12:12 hour light:dark cycle. Following jugular catheter implantation, mice were singly housed and maintained on a 14:10 hour light:dark cycle. Experiments were conducted during the light period, with the exception of extended access self-administration training, which took place predominantly during the dark period. To control for circadian effects (*55, 56*), the time of self-administration was held constant for individual animals throughout the study. Tap water and standard laboratory chow were supplied *ad libitum*, except during testing. Studies were conducted with 3 and 2 replicate cohorts of animals, for cocaine and remifentanil, respectively. Replicate cohorts were run sequentially. All mouse studies were conducted in accordance with the National Institutes of Health Guidelines for Animal Care and Use and with approved animal protocols from the Duke University Institutional Animal Care and Use Committee.

### Drug treatment

Cocaine hydrochloride (cocaine) was purchased from MilliporeSigma (St. Louis, MO) and remifentanil hydrochloride (remifentanil) was provided by the National Institute on Drug Abuse (NIDA), through the NIDA Drug Supply Program. Both drugs were dissolved in sterile physiological saline (Henry Schein, Raleigh, NC). For open field studies, cocaine (20 mg/kg, ip) or remifentanil (0.01 – 10 mg/kg, ip) was injected in a volume of 10 ml/kg body weight. For iv self-administration studies, cocaine (0.1-3 mg/kg/infusion) or remifentanil (0.01-3 mg/kg/infusion) was administered intravenously in a volume of 18 µl/infusion to animals with indwelling jugular catheters according to the test protocol and their lever responding.

### Jugular catheterization

Indwelling catheters (Instech Laboratories, Inc., Plymouth Meeting, PA) were implanted into the right jugular vein of adult male and female C57BL/6J mice, as described for CD-1 mice (*41*), with several modifications. Specifically, changes were made in the type and route of anesthesia, the design/supplier of the catheters and vascular access buttons, the material of the vascular access buttons, and the size of the insertion needle. Briefly, mice were anaesthetized with a freshly prepared ketamine/xylazine (12/1 mg/ml) cocktail. Catheters were inserted into the right jugular vein and threaded under the skin of the shoulder to a mid-scapular vascular access button. Polyurethane catheters had a bead 1.2 cm from the catheter tip (Instech, Plymouth Meeting, PA; Part #VAB62BS/25). To prepare the catheters for insertion, the excess catheter tubing was cut 3.5 cm from the bead and affixed to a vascular access button (Instech, Plymouth Meeting, PA; Part #KVAH62T). A modified 18-gauge, 1.5 inch guide needle (Becton Dickinson, Franklin Lakes, NJ; Part # BD-305196) was used for catheter insertion. On post-surgery days 1-3, animals received amikacin at a dose of 10 mg/kg (s.c.) to prevent perioperative infection. To maintain patency and prevent infection for the duration of the study, catheters were flushed 1-2x daily with 50 μl of a heparin/gentamicin solution.

### Jugular catheter patency testing

The patency and optimal placement of jugular catheters was determined by infusing 20-30 μl of a 15 mg/ml ketamine solution in sterile saline (*41, 57*). In this test, the ketamine infusion resulted in rapid sedation (i.e., within 3 sec), as evidenced by loss of muscle tone in animals with patent catheters. Catheter malfunctions were manifested as a delayed time to onset of sedation (i.e., >3 sec) (*57, 58*) or an inability to infuse the ketamine; these mice were excluded from the study. Animals that showed evidence of sedation within 3 sec of infusion were considered to have patent catheters. Patency testing was conducted with all mice on post-surgery day 7 to verify successful catheter placement, as well as according to behavioral performance criteria throughout the experiment, and at the completion of the study. Behavioral performance that qualified animals for catheter patency testing included a failure to progress toward training criteria after 4 consecutive sessions on the same protocol, loss of the active lever preference in trained animals, and/or inconsistencies in lever responding that were not concurrent with changes in the animal’s body weight, nesting activity or activity level.

### Locomotor activity

The VersaMax activity monitor (Omnitech Electronics, Inc.) was used to assess locomotor activity, as previously described (*59*). Male and female mice with indwelling jugular catheters were acclimated to the open field for 30 min prior to administration (ip) of cocaine (20 mg/kg, 10 ml/kg) or remifentanil (0.1, 1.0 and 10 mg/kg, 10 ml/kg, sequentially). After cocaine treatment, animals were immediately returned to the open field and locomotion was monitored over the next 90 min. After each successive remifentanil treatment, animals were immediately returned to the open field and activity was monitored over the next 20 min.

### Self-Administration

See **Figures S1 and S2** for protocol summaries.

#### Acquisition

Following a 7-14 day post-surgical recovery period, mice that completed *a priori* open field assessments and passed a ketamine test of catheter patency (*41*) were trained to iv self-administer cocaine (0.5 mg/kg/infusion) or remifentanil (0.1 mg/kg/infusion) paired with a cue light through lever responding in operant chambers (Med Associates, Inc., Fairfax, VT). Active, reinforced lever(s) were denoted by a solid cue light. Reinforcements were delivered by a single speed syringe pump (Med Associates) in a volume of 18 μl over a 4 sec period. Each infusion was followed by a 40-sec time-out period during which no additional reinforcements were provided. This time-out period was imposed to prevent adverse health consequences associated with excess cocaine or remifentanil administration (*60*). For the first 20 sec of this time-out period, the stimulus light blinked and the levers remained available, but lever responses (i.e., time-out responses) had no programed consequences. For the following 20 sec, the levers were retracted. Training began with an autoshaping session followed by a 12 hr extended access session in which responding on either of two levers resulted in reinforcement. Subsequently, in 1 hr daily sessions, mice were trained to discriminate an active drug-delivering lever from an inactive non-drug-delivering lever and the fixed lever response to reinforcement ratio (i.e., fixed ratio, FR) was increased from FR1 to FR2 to FR4. Training followed the contingent advancement protocols provided (**Figure S1, S2**). A small control group was given access to the vehicle (saline) rather than cocaine or remifentanil. This group completed a single autoshaping and a single extended-access session followed by 5 consecutive sessions of each training protocol.

#### Maintenance dose-response testing

After drug self-administration was acquired, stable lever responding at FR4 was assessed at 5 cocaine and 6 remifentanil doses. For cocaine, doses (mg/kg/infusion) were presented as: 0.5, 0.3. 0.1, 1.0, 3.0. The remifentanil, doses (mg/kg/infusion) were presented in the following order: 0.1, 0.03, 0.01, 0.3, 1.0, 3.0. The 3.0 cocaine and remifentanil doses were added after the first replicate cohort had completed the study to better define the descending portion of the drug dose-effect curves. Stable responding was considered to have been achieved when self-administered reinforcements in two consecutive sessions varied by ≤ 20% with ≥ 50% of the total responses occurring at the active lever. Animals that failed to achieve or maintain stable responding and ≥ 50% active lever discrimination over 4 consecutive sessions were subjected to a patency test. Non-patent animals were removed from the study.

#### Extinction

After completing maintenance dose-response testing, mice were subjected to once daily, 1-hour sessions in which the cue light was absent and lever responses were recorded but they had no programmed consequences. For the cocaine paradigm, daily extinction sessions continued until lever responses at the animal’s previously active lever were reduced to ≤ 20% of the animal’s previous responses for the 1 mg/kg/infusion cocaine dose over 2 consecutive sessions or until 40 total sessions were completed. Because preliminary cocaine data suggested the rate of extinction plateaued by session 20, in the remifentanil paradigm, all animals completed 20 daily extinction sessions regardless of performance.

#### Cue-induced reinstatement

After meeting extinction criteria, mice completed a single cue-induced reinstatement session. In this session, the drug-associated light was illuminated over the previously active lever and responses on this lever resulted in presentation of other drug-associated cues, including the cue light blinking, lever movements, and the sound of the syringe pump delivery, in the absence of drug reinforcement.

### Data capture and storage

Self-administration data were collected and managed in a customized REDCap (Research Electronic Data Capture) database created using REDCap electronic data capture tools hosted at Duke University (*61, 62*). Note, REDCap is a secure, clinically-utilized web-based software designed to support data capture for research studies.

### Statistical analysis

All data are represented as mean ± SEMs, unless otherwise indicated. Primary behavioral data were analyzed and plotted using the software GraphPad Prism version 8.0. Information on curve fitting and statistical assessments are provided in the text, figure legends or supplemental tables. Statistical significance was assigned at p<0.05.

#### Correlation matrices and hierarchical clustering

Pearson correlation coefficients for self-administration variables and novelty- and drug-induced activity open field variables were determined using the ‘cor’ function in R-Studio version 3. Agglomerative hierarchical clustering and dendrograms for self-administration variables were built using the R-Studio’s ‘hclust’ function with complete-linkage.

#### Exploratory factor analysis

To investigate the potential dimensional inter-relationships between self-administration behaviors at the stages of acquisition, maintenance, extinction, and cue-induced reinstatement, within-drug, correlation matrices and R-type exploratory factor analyses (EFAs) were performed using the IBM SPSS Statistics version 26 program. A total of 12 variables from the cocaine self-administration paradigm and 11 variables from the remifentanil self-administration paradigm were included in the models. Linear relationships were visually confirmed using bivariate scatterplots. All variables remained on their natural scales and the robustness of the models to the assumption of multivariate normality was assessed. Missing values were replaced with the within-drug variable mean. Although 4 factors with eigen values >1.0 were identified in the cocaine and remifentanil models, because the objective was dimension reduction, we extracted by maximum likelihood estimation only the two factors with the highest explanatory values. Unrotated loading plots and factor matrices were generated. Extracted factor 1 and extracted factor 2 scores were calculated for individual mice by regression. Only animals that met acquisition criteria and achieved stable responding at FR4 with 0.5 mg/kg/infusion of cocaine or 0.1 mg/kg/infusion of remifentanil were included in the models (n=38 cocaine; n=34 remifentanil), yielding a case to variable ratio of ∼3:1.

To determine whether cocaine and remifentanil reinforcement produced distinct global self-administration phenotypes, a third factor analysis was performed that included data from 11 common variables with all mice irrespective of reinforcer (n = 72). Missing values were replaced with the global variable mean. The two factors with the highest eigen values were again extracted, by maximum likelihood estimation. Extracted factor 1 and extracted factor 2 scores from the unrotated solution were calculated for individual animals by regression. Factor 1 and 2 scores were plotted and compared by reinforcer.

To assess the relationship between behavior in the open field and self-administration parameters, Pearson’s R correlation coefficients were calculated between novelty- and drug-induced open field activity measures and within-drug factor scores (n= 28 cocaine; n= 34 remifentanil). Since open field data were not collected from an initial set of 10 mice in the cocaine self-administration paradigm, they were excluded from the analysis.

#### Predictive modeling of drug seeking

Machine learning was used to develop predictive models of early extinction, late extinction and reinstatement lever responding based on novelty-induced responses in the open field, sensitivity to drug-induced hyperlocomotion and drug taking behavior. The data were split into a training and a test dataset using a random 80:20 split and models were built based on the training dataset. To remove multicollinearity in the novelty open field and drug-taking data, principle component analysis (PCA) was performed in R-Studio version 3. Prior the PCA, data for 6 open field variables and 8 drug-taking-associated variables were first scaled using the scikit-learn package for Python. The number of components extracted was determined by the minimum required to cumulatively explain 97% of the variance. Extracted components and drug-induced locomotion data were treated as features to build multiple linear regressions using the ‘lm’ function in R-Studio. The ‘step’ function was used to select independent variables based on Akaike information criterion (*63, 64*). The trained models were applied to test datasets to make predictions. Mean absolute errors from the training and test datasets were calculated and compared to assess the model performance.

### Figure illustration

The study overview figures presented in Figure 1 was created using BioRender (Toronto, ON).

## SUPPLEMENTAL INFORMATION

Attached as a separate file.

## ACKNOWLEDGEMENTS

We thank Xiuqin Zhang and Wendy L. Roberts for managing our mouse colony, and Dr. Ramona M. Rodriguiz for coordinating experimental schedules. This work was supported, in part, by NIH/NIDA grants P30 DA029925, F32 DA043931, and K99 DA048970.

## AUTHOR CONTRIBUTIONS

Study design: LMS, AP, YB, NC, VMP, KT, WCW, LSB and MGC. Obtained funding: LMS, LSB, MGC. Data collection: LMS, AP, YB, NC, KT and FP. Data analysis: LMS, AP, NC, FP, EH, YZ and XC. Data interpretation: LMS, ERH, VMP, KT, JDG, WCW, LSB and MGC. Drafted paper: LMS. Revised paper critically for intellectual content: LMS, ERH, JDG, VMP, WCW, LSB and MGC.

## DECLARATION OF INTERESTS

The authors declare no competing interests.

## Notes

### Competing Interest Statement

The authors have declared no competing interest.

### Summary of Updates

Figure 1 added; Figure 5 revised; Figure 6 revised; Author list updated; Supplemental files updated

