## Supplementary Material for "Establishment of Multi-stage Intravenous Self-administration Paradigms in Mice"

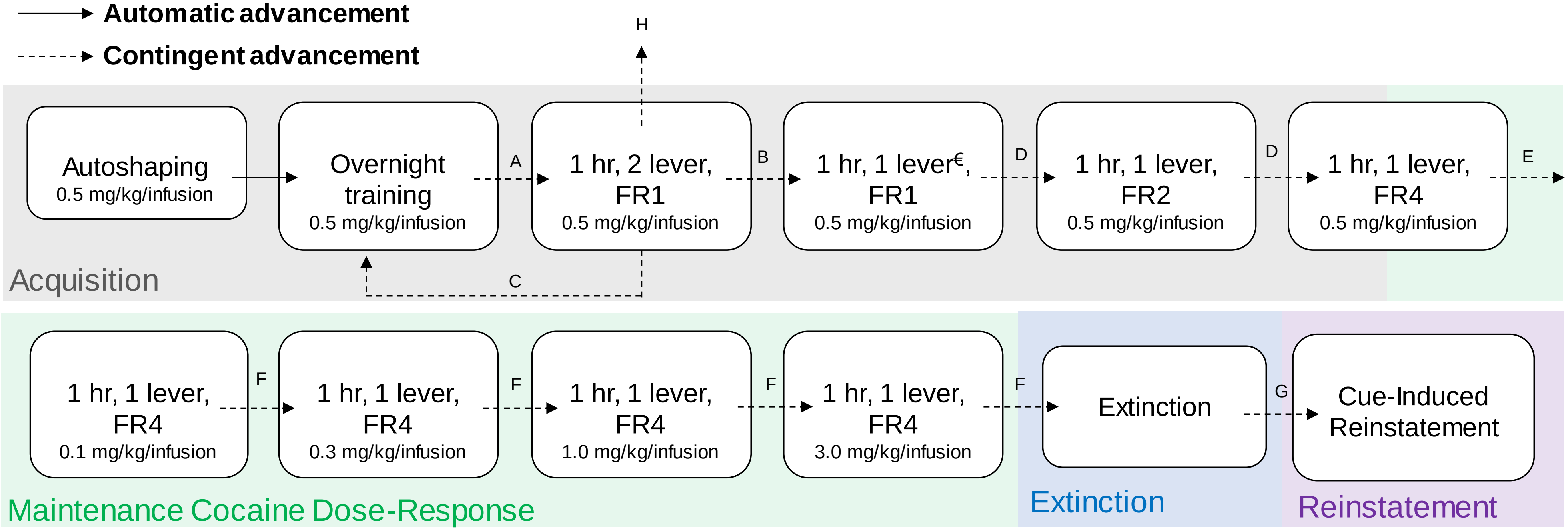


**Figure S1. Cocaine self-administration training and testing paradigm.** Animals were trained to self-administer i.v. cocaine paired with a cue light via lever pressing in operant chambers. A schematic of the training and testing protocol is presented above. *Solid arrows* denote phases between which animals progressed automatically. *Dashed arrows* indicate that progression to the following phase was contingent on meeting predetermined behavioral criteria. Advancement criteria were as follows: **A.** ≥ 50 rewards; **B.** ≥ 10 rewards in 2 consecutive sessions; **C.** ≤ 9 rewards in 3 consecutive sessions; **D.** ≥ 10 rewards with ≥ 50% presses on the active lever in 2 consecutive sessions; **E.** ≥ 2 sessions with ≥ 10 rewards, ≥ 50% presses on the active lever and ≤ 20% variability in rewards in the last 2 consecutive sessions *OR* ≥ 3 sessions with <10 rewards, ≥ 50% presses on the active lever and ≤ 20% variability in rewards in the last 2 consecutive sessions; **F.** ≥ 3 sessions with ≥ 50% presses on the active lever and ≤ 20% variability in rewards in the last 2 consecutive sessions; **G.** ≤ 20% of active lever presses in the last cocaine self-administration session; **H.** Animals not meeting self-administration criteria after 4 overnight training sessions and 12 1 hr, 2 lever, FR1 training sessions were terminated from study. [€]The lever selected as active was the non-preferred lever during last two lever training session. Animals not meeting criteria for advancement repeated their current protocol until criteria were met. Failure to progress toward criteria after 4 consecutive sessions on the same protocol prompted a ketamine test for catheter patency.

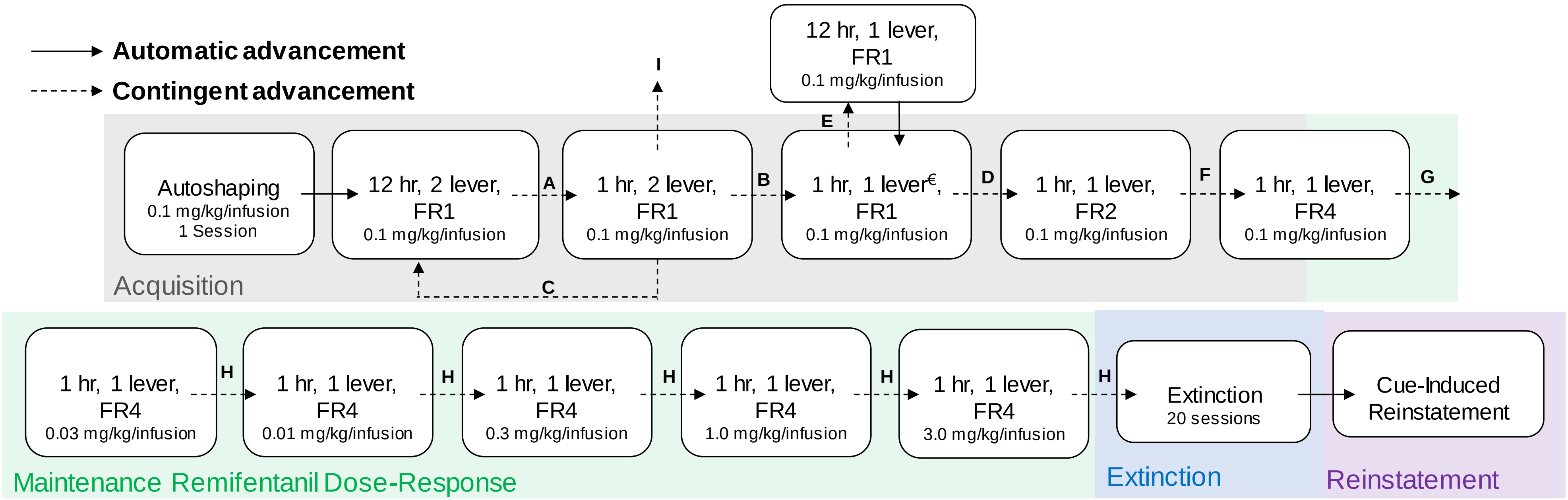


**Figure S2. Remifentanil self-administration training and testing paradigm.** Animals were trained to self-administer i.v. remifentanil paired with a cue light via lever pressing in operant chambers. A schematic of the training and testing protocol is displayed above. *Solid arrows* denote phases between which animals progressed automatically. *Dashed arrows* indicate that progression to the following phase was contingent on meeting predetermined behavioral criteria. Advancement criteria were as follows: **A.** ≥ 50 rewards; **B.** ≥ 10 rewards in 2 consecutive sessions; **C.** ≤ 9 rewards in 3 consecutive sessions; **D.** ≥ 10 rewards with ≥ 50% presses on the active lever in 2 consecutive sessions; **E.** <10 rewards or <50% presses on the active lever in 6 consecutive sessions; **F.** ≥ 10 rewards, ≥ 50% presses on the active lever in 2 consecutive sessions or in ≥ 3 sessions with <10 rewards, ≥ 50% presses on the active lever and ≤ 20% variability in rewards in the last 2 consecutive sessions; **G.** ≥ 2 sessions with ≥ 10 rewards, ≥ 50% presses on the active lever and ≤ 20% variability in rewards in the last 2 consecutive sessions *OR* ≥ 3 sessions with <10 rewards, ≥ 50% presses on the active lever and ≤ 20% variability in rewards in the last 2 consecutive sessions; **H.** ≥ 3 sessions with ≥ 50% presses on the active lever and ≤ 20% variability in rewards in the last 2 consecutive sessions; **I.** Animals not meeting self-administration criteria after 4 overnight training sessions and 12 1 hr, 2 lever, FR1 training sessions were excluded from study. €The lever selected as active was the non-preferred lever during last two lever training session. Animals not meeting criteria for advancement repeated their current protocol until criteria were met. Failure to progress toward criteria after 4 consecutive sessions on the same protocol prompted a ketamine test for catheter patency.

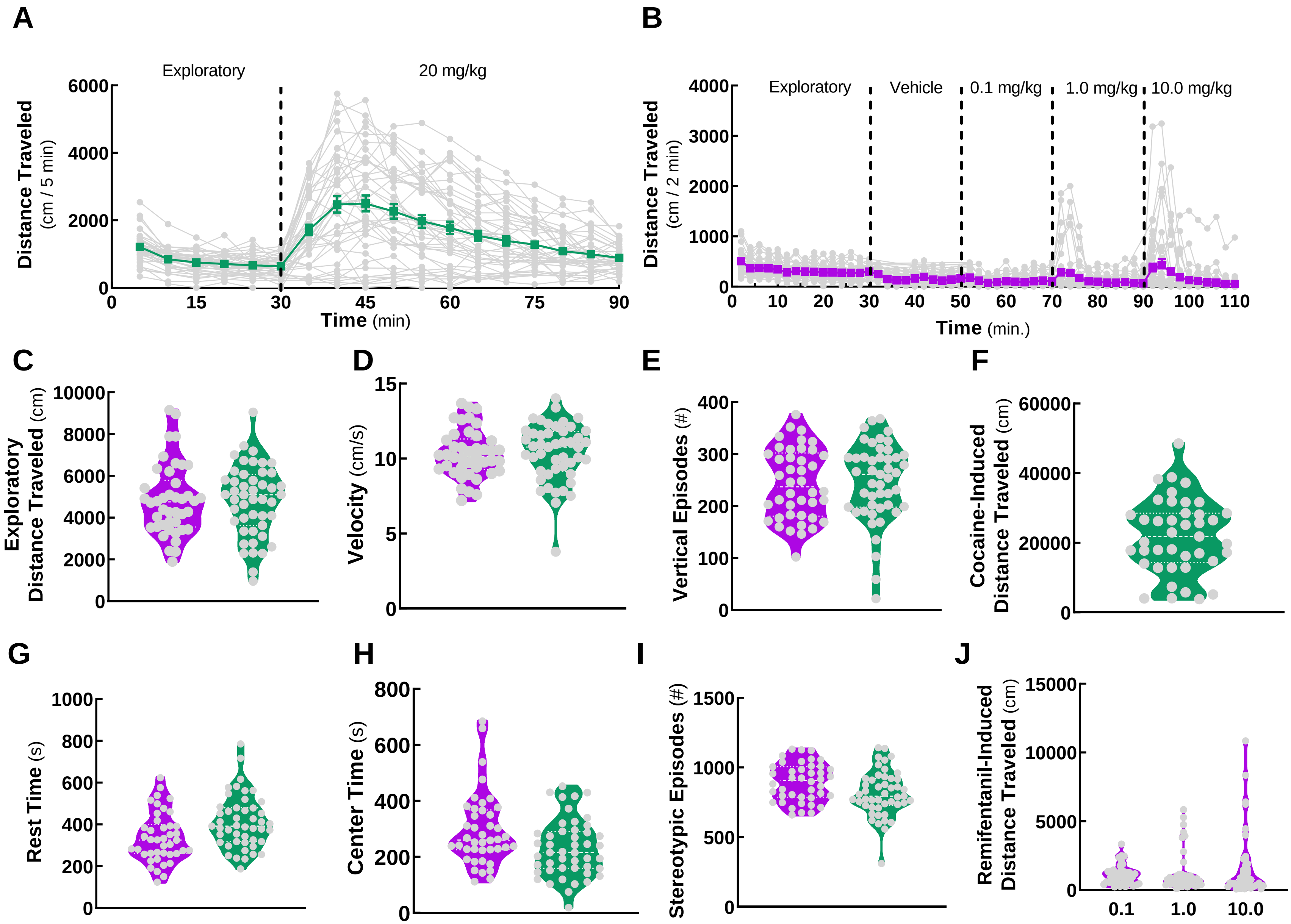
A
Exploratory
20 mg/kg
Distance Traveled (cm / 5 min)
Time (min)
B
Exploratory
Vehicle
0.1 mg/kg
1.0 mg/kg
10.0 mg/kg
Distance Traveled (cm / 2 min)
Time (min.)
C
Exploratory Distance Traveled (cm)
D
Velocity (cm/s)
E
Vertical Episodes (#)
F
Cocaine-Induced Distance Traveled (cm)
G
Rest Time (s)
H
Center Time (s)
I
Stereotypic Episodes (#)
J
Remifentanil-Induced Distance Traveled (cm)
0.1
1.0
10.0

**Figure S3. Exploratory and cocaine- and remifentanil-induced locomotor activities.** Exploratory and cocaine- or remifentanil-induced locomotor activities were evaluated in the open field using automated activity monitors. Mice were acclimated to the open field for 30 min prior to drug administration. Cocaine (20 mg/kg) or vehicle (saline) was injected (i.p.) and the mice were returned immediately to the open field for 60 min. After collecting baseline activity, the mice to receive remifentanil were injected (i.p.) sequentially with the vehicle (saline), followed in 20 min inervals by consecutive remifentanil treatments at doses of 0.1, 1.0 and 10.0 mg/kg (i.p.). Data are presented as mean ± SEM in panels **(A)** and **(B)**; group means are represented by *green* (cocaine) or *purple* (remifentanil) *lines* and individual replicates by *gray lines*. Panels **(C-J)** are displayed as violin plots with individual data points represented by *grey circles*. Remifentanil data are shown in *purple* and cocaine data are depicted in *green*.

**(A) Time-course of exploratory and cocaine-induced locomotion.** Distance traveled (cm) in 5-min bins over the 60 min post-treatment period.

**(B) Time-course of exploratory and remifentanil-induced locomotion.** Distance traveled (cm) in 2-min bins over 20 min successive post-sequential treatments.

**(C) Cumulative exploratory distance traveled.**

**(D) Mean exploratory velocity.**

**(E) Cumulative exploratory vertical episodes.**

**(F) Cumulative cocaine-induced distance traveled.**

**(G) Cumulative exploratory rest time.**

**(H) Cumulative exploratory center time.**

**(I) Cumulative exploratory stereotypic episodes.**

**(J) Cumulative remifentanil-induced distance traveled.**

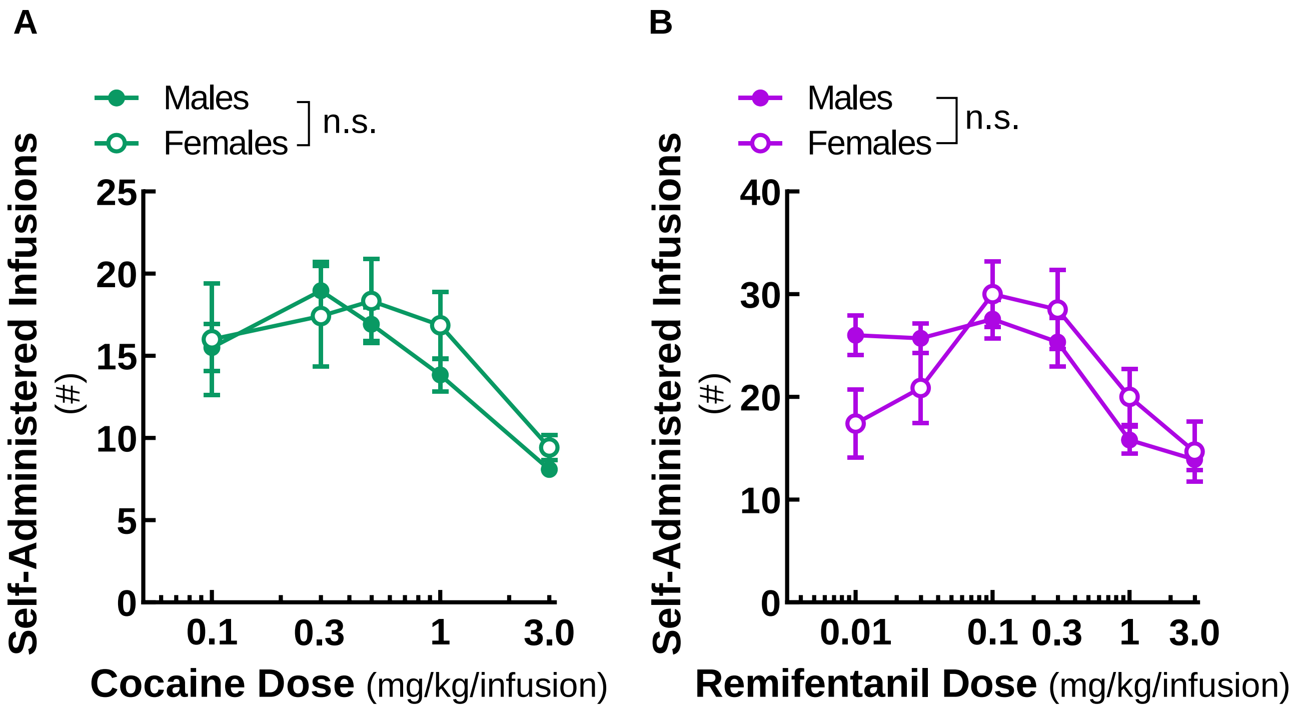


**Figure S4. Reinforcement dose-response curve by sex.** No effect of sex on self-administered **(A)** cocaine or **(B)** remifentanil reinforcements were identified. Data were analyzed using two-way, mixed model ANOVA. Cocaine: n = 20-28 males, 7-9 females. $F_{Dose}(2.95, 88.39)=18.1$, $p<0.0001$; $F_{Sex}(1, 36)=0.0004$, $p=0.9844$; $F_{Interaction}(4, 120)=1.1$, $p=0.3588$. Remifentanil: n = 12-26 males, 3-8 females. $F_{Dose}(3.37, 85.03)=8.89$, $p<0.0001$; $F_{Sex}(1, 32)=0.000929$, $p=0.9759$; $F_{Interaction}(5, 126)=2.48$, $p= 0.0352$. n.s., not significant**.**

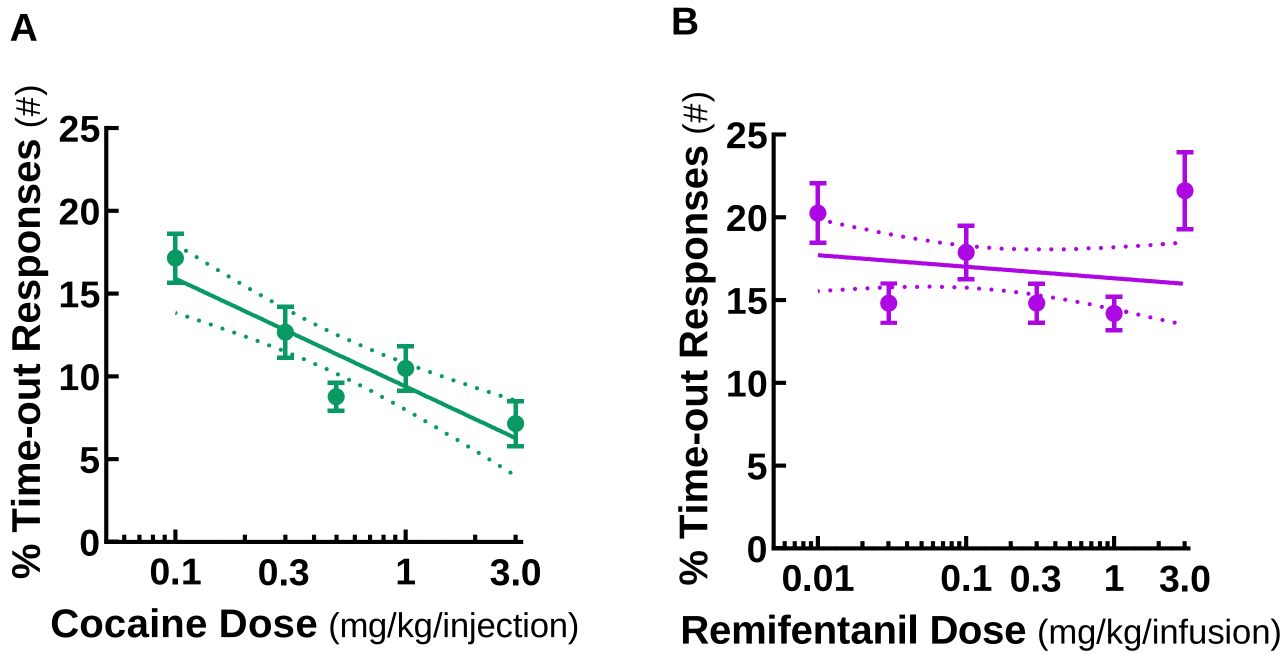


**Figure S5. Percent of total lever responses that occurred during the post-reinforcement time-out period by drug dose.** Percent time-out lever responses were fit to linear regressions (*solid lines*) with the 95% confidence limits of the best-fit line represented by *dashed lines*. **(A)** In the cocaine paradigm, the percent of total lever responses that occurred during the time-out period decreased linearly with log cocaine dose. Y=-6.5X + 9.4. Slope 95% confidence interval: -9.0 to -4.1 time-out responses per log mg/kg/infusion cocaine. **(B)** In the remifentanil paradigm, the percent of total lever responses that occurred during the time-out period was unrelated to the log remifentanil dose. Y=-.07X + 16.3. Slope 95% confidence interval: -2.2 to 0.9 time-out responses per log mg/kg/infusion remifentanil.

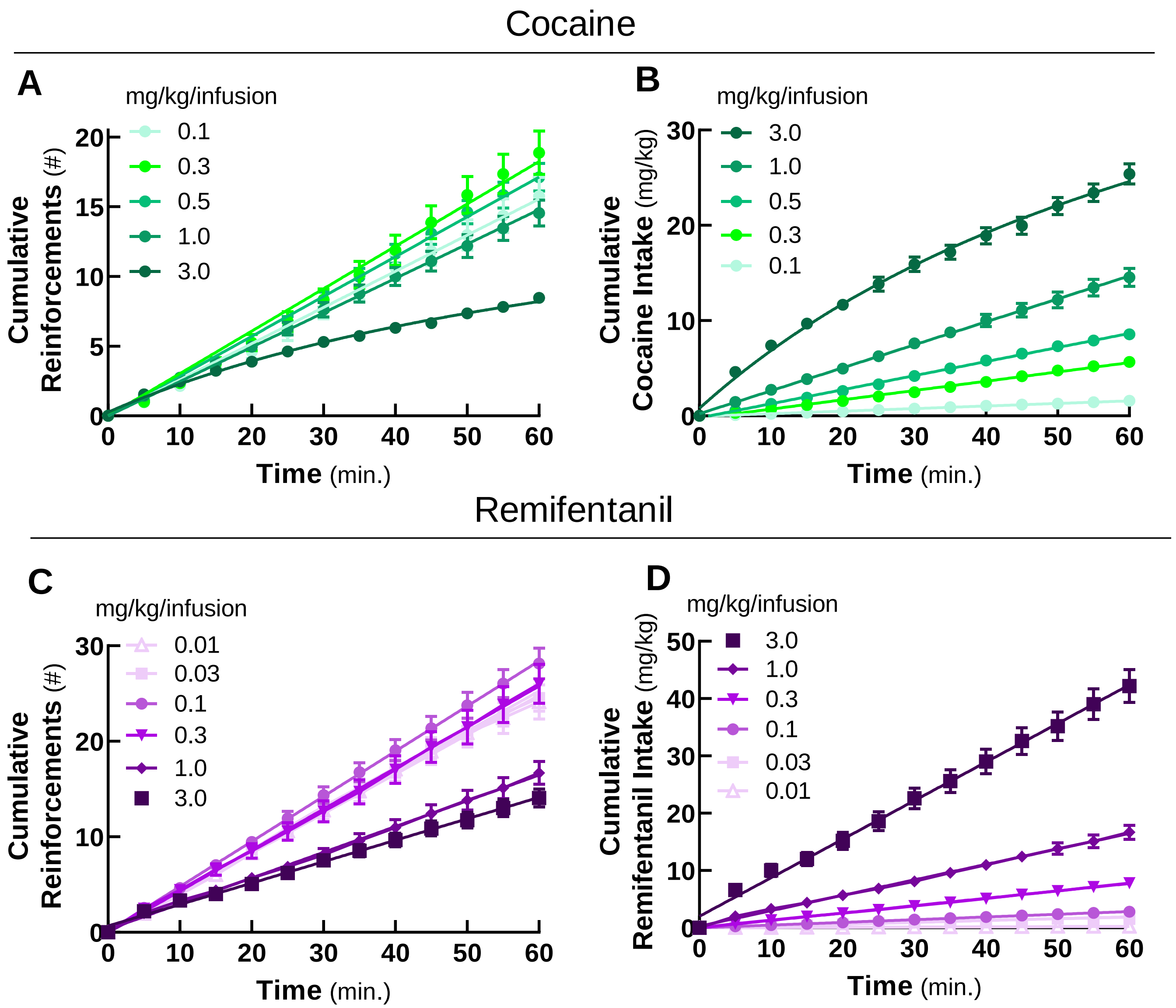


**Figure S6. Kinetics of self-administration over 60 min sessions.** Cumulative self-administered reinforcements for cocaine (0.1-3.0 mg/kg/infusion, **A**) and remifentanil (0.01-3.0 mg/kg/infusion, **C**) over 60 min self-administration sessions. Cumulative drug intake for cocaine (0.1-3.0 mg/kg/infusion, **B**) and remifentanil (0.01-3.0 mg/kg/infusion, **D**) over 60 min self-administration sessions. Data for all remifentanil doses were fit by linear regression. Data for cocaine doses at and below 1.0 mg/kg/infusion were fit by linear regression. The 3.0 mg/kg/infusion cocaine dose curves were fit to rectangular hyperbolas.

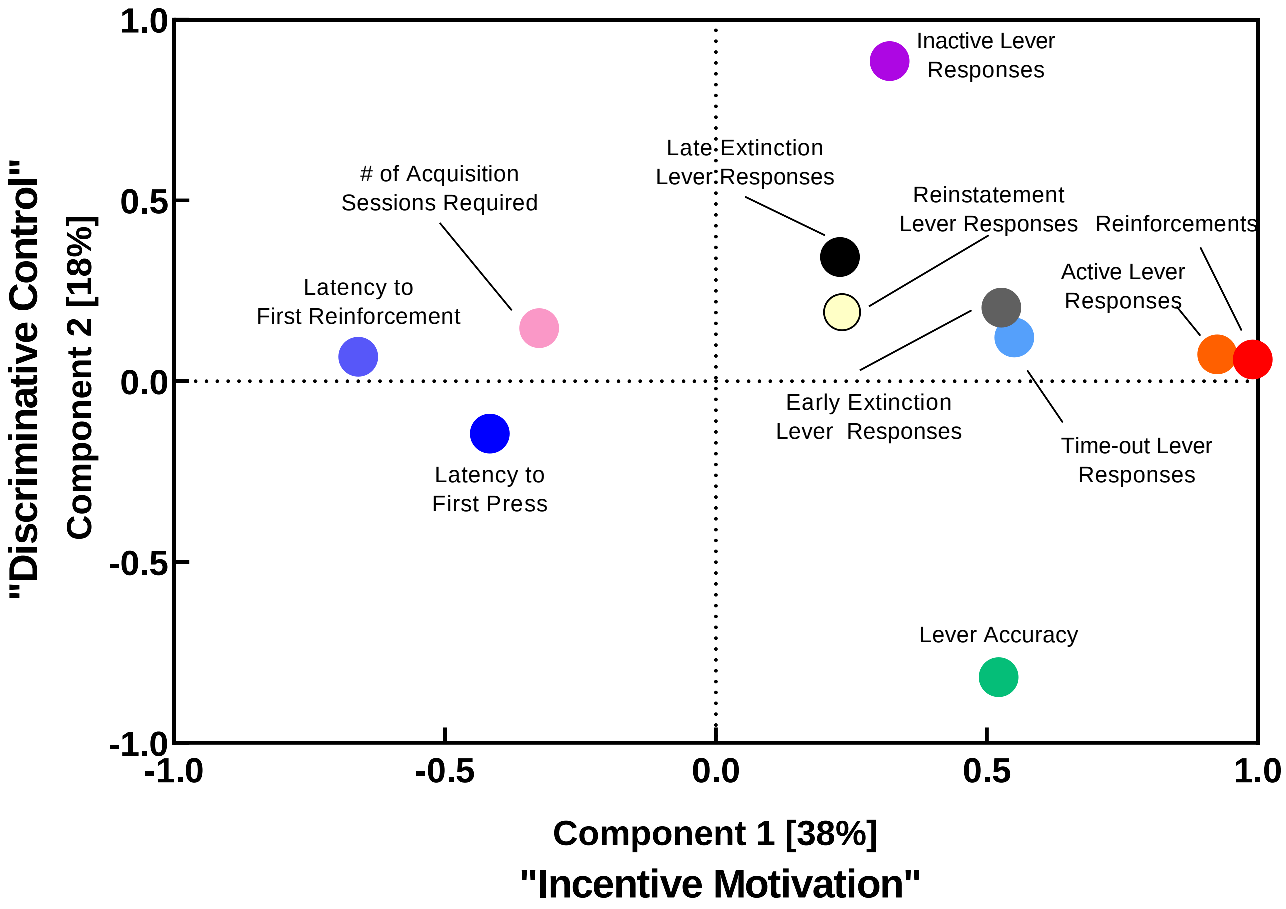


**Figure S7. Factor loading plot of variables in global factor analysis.** Loadings of 11 observed variables included in the global exploratory factor analysis on extracted factors 1 and 2 are presented. The term global is used to indicate that all animals were included in the analysis, regardless of the reinforcer/drug paradigm. Variable loading scores are included in **Table S4**.

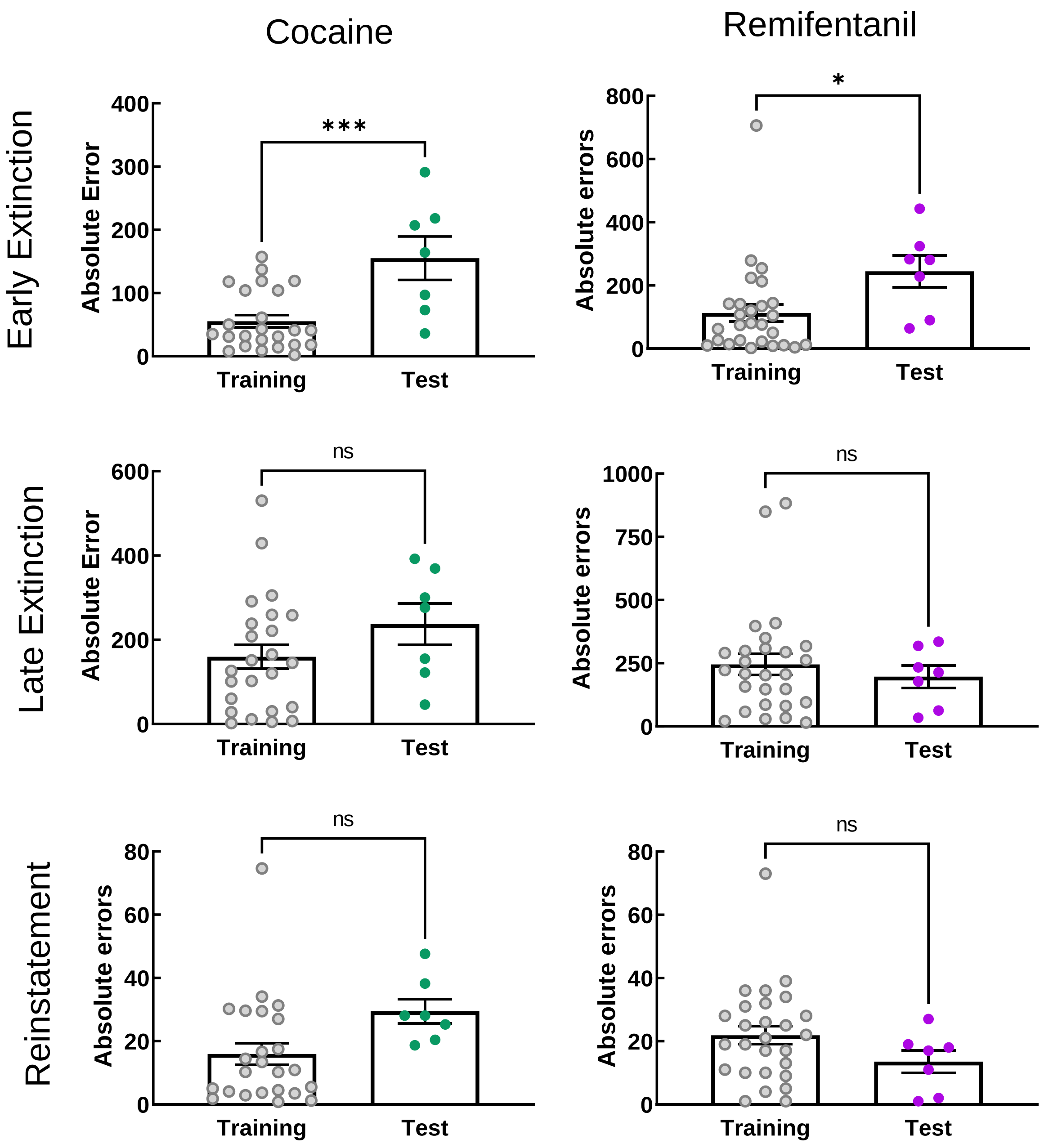


**Figure S8. Machine learning regression model performance.** Absolute residual values from actual vs. predicted plots in **Figure 5**. Mean absolute error values between training and test sets were compared using *Student's t-tests*.

**Table S1. Self-administration acquisition – Supporting Figure 2.**

| | Cocaine | Remifentanil | P value* |
|---|---|---|---|
| Sessions to meet extended access criteria | 1.2 ± 0.4 | 1.1 ± 0.4 | 0.0931 |
| Sessions to meet 2 lever, FR1 criteria | 3.3 ± 2.0 | 2.2 ± 0.4 | 0.0010 |
| Sessions to meet 1 lever, FR1 criteria | 4.0 ± 1.6 | 2.9 ± 0.8 | 0.0009 |
| Sessions to meet 1 lever, FR2 criteria | 2.1 ± 0.5 | 2.0 ± 0.2 | 0.4767 |
| Sessions to meet 1 lever, FR4 criteria | 2.8 ± 1.6 | 2.6 ± 0.8 | 0.7891 |
| Total required training sessions | 13.5 ± 3.8 | 10.8 ± 1.2 | 0.0002 |
| Mice trained (n) | 38 | 34 | |
| Mice failed to train (n) | 7 | 2 | |
| Percent mice completing training | 84.4 | 94.4 | 0.1547[¥] |

Mean number of sessions required to meet specific criteria ±SD. *Mann-Whitney test. ¥Chi-square test.

### Table S2. Statistical analysis of data presented in Figure 1 – Supporting Figure 2.

| Figure | Panel | Experiment Description | F Statistics[1] | Multiple Comparisons[2] | N[3] | Curve Fit | Parameter | 95% CI | Units |
|---|---|---|---|---|---|---|---|---|---|
| 1 | A | Number and proportion of mice that met final SA acquisition criteria, **Cocaine** | | | 42 | | | | |
| | B | Number of sessions required to meet training criteria, **Cocaine** | | | 37 | Nonlinear regression, Lorentizian fit | Center<br>Width<br>$R^2$<br><br>Mean<br>Median<br>SD | 8.1 to 12.7<br>0.9 to 7.7<br>0.6093<br><br>13.5<br>12.0<br>3.8 | sessions<br>sessions<br><br><br>sessions<br>sessions |
| | C | Number and proportion of mice that met final SA acquisition criteria, **Remifentanil** | | | 36 | | | | |
| | D | Number of sessions required to meet training criteria, **Remifentanil** | | | 34 | Nonlinear regression, Lorentizian fit | Center<br>Width<br>$R^2$<br><br>Mean<br>Median<br>SD | 10.0 to<br>11.1<br>-2.3 to 2.3<br>0.9060<br><br>10.8<br>11<br>1.3 | sessions<br>sessions<br><br><br>sessions<br>sessions |
| | E | Lever responses vs. training session type, **Cocaine** | $F_{Session}(3.34, 200.2)=23.7, p<0.0001$<br>$F_{Lever}(1,74)=32.4, p<0.0001$<br>$F_{Interaction}(8, 480)=49.7, p<0.0001$<br><br>Geisser-Greenhouse's ε=0.42 | ***p<0.0001, Active vs. Inactive Lever | 38 | | | | |
| | F | Lever responses vs. training session type, **Remifentanil** | $F_{Session}(4.35, 208.4)=15.1, p<0.0001$<br>$F_{Lever}(1,66)=44.3, p<0.0001$<br>$F_{Interaction}(8, 383)=33.9, p<0.0001$<br><br>Geisser-Greenhouse's ε=0.54 | ***p<0.0001, Active vs. Inactive Lever | 34 | | | | |
| | G | Active lever responses vs. FR, **Cocaine** | F(1.55, 55.87)=71.0, p<0.0001<br><br>Geisser-Greenhouse's ε=0.89 | ***p<0.001, FR1 vs. FR2, FR1 vs. FR4, FR2 vs. FR4 | 38 | | | | |
| | H | Inactive lever responses vs. FR, **Cocaine** | F(1.80, 64.51)=6.234, p=0.0045<br><br>Geisser-Greenhouse's ε=0.90 | *p<0.05, FR1 vs. FR2<br>*p<0.01, FR1 vs. FR4 | 38 | | | | |

| | | | | |
|---|---|---|---|---|
| **I** | Active lever responses vs. FR,<br>**Remifentanil** | F(1.80, 57.65)=33.3, p<0.0001<br><br>Geisser-Greenhouse's ε=0.90 | *p<0.01, FR1 vs. FR2<br>***p<0.0001, FR1 vs. FR4, FR2 vs. FR4 | 34 |
| **J** | Inactive lever responses vs. FR,<br>**Remifentanil** | F(1.945, 62.23)=0.7, p=0.5019<br><br>Geisser-Greenhouse's ε=0.97 | n.s. | 34 |
| **K** | Lever accuracy vs. FR,<br>**Cocaine** | F(1.95, 70.20)=5.4, p=0.0072<br><br>Geisser-Greenhouse's ε=0.98 | *p<0.01, FR1 vs. FR4 | 38 |
| **L** | Reinforcements vs. FR,<br>**Cocaine** | F(1.77, 63.77)=20.5, p<0.0001<br><br>Geisser-Greenhouse's ε=0.89 | *p<0.05, FR1 vs. FR2<br>***p<0.001, FR1 vs. FR4<br>**p<0.001 FR2 vs. FR4 | 38 |
| **M** | Lever accuracy vs. FR,<br>**Remifentanil** | F(1.97, 63.09)=5.6, p=0.0057<br><br>Geisser-Greenhouse's ε=0.96 | *p<0.01 FR1 vs. FR4 | 34 |
| **N** | Reinforcements vs. FR,<br>**Remifentanil** | F(1.94, 62.03)=35.5, p<0.0001<br><br>Geisser-Greenhouse's ε=0.97 | *p<0.05, FR1 vs. FR2<br>***p<0.0001, FR1 vs. FR4, FR2 vs. FR4 | 34 |

[1]F statistics from two-way, mixed effects analyses for panels E and F and one-way, repeated measures ANOVAs for panels G-L; [2]*Post-hoc* Sidak tests for panels E and F and Tukey's multiple comparisons tests for panels G-L. [3]A single mouse in each of the cocaine and remifentanil paradigms was removed from training analyses because they were prematurely progressed to FR4 due to experimenter error. Sphericity was not assumed, and the analyses were corrected using the Geisser-Greenhouse epsilon (ε) hat method; CI, confidence internal; SD, standard deviation; n.s., not significant.

#### Table S3. Curve parameters presented in Figure 2 – Supporting Figure 3.

| Figure | Panel | Experiment Description | Curve Fit | N | Parameter | 95% CI | Units |
|---|---|---|---|---|---|---|---|
| **2** | **A** | Earned reinforcements vs. log cocaine dose | Nonlinear regression, second order polynomial | 27-38 | B0<br>B1<br>B2<br>$R^2$ | 13.8 to 16.5<br>-13.3 to -7.1<br>13.6 to -5.4<br>0.2042 | mg/kg/infusion cocaine<br>mg/kg/infusion cocaine<br>mg/kg/infusion cocaine |
| | **B** | Total active lever responses vs. log cocaine dose | Nonlinear regression, second order polynomial | 27-38 | B0<br>B1<br>B2<br>$R^2$ | 58.8 to 68.7<br>56.6 to -33.0<br>-40.6 to -10.1<br>0.1481 | mg/kg/infusion cocaine<br>mg/kg/infusion cocaine<br>mg/kg/infusion cocaine |
| | **C** | Total inactive lever responses vs. log cocaine dose | Linear regression | 27-38 | Slope<br>Y-intercept<br>$R^2$ | 15.9 to -10.1<br>9.34 to 12.66<br>0.1644 | lever response per mg/kg/infusion cocaine<br>lever responses |
| | **D** | Cocaine consumed vs. log cocaine dose | Nonlinear regression, sigmoidal | 27-38 | Bottom<br>Top<br>$EC_{50}$<br>Hill Slope<br>$R^2$ | Constrained to 0<br>30.6 to 61.8<br>1.0 to 4.7<br>0.8 to 1.3<br>0.7704 | <br>mg/kg cocaine<br>mg/kg/infusion cocaine |
| | **E** | Percent active lever responses vs. log cocaine dose | Linear regression | 27-38 | Slope<br>Y-intercept<br>$R^2$ | 7.5 to 14.4<br>84.9 to 88.8<br>0.1942 | percent active lever responses per mg/kg/infusion cocaine<br>percent active lever responses |
| | **F** | Latency to first lever response vs. log cocaine dose | Linear regression | 27-38 | Slope<br>Y-intercept<br>$R^2$ | -32.9 to 14.89<br>27.9 to 55.2<br>0.003387 | sec per mg/kg/infusion cocaine<br>sec |
| | **G** | Latency to first earned reinforcement vs. log cocaine dose | Linear regression | 27-38 | Slope<br>Y-intercept<br>$R^2$ | 174.1 to -54.98<br>149.9 to 217.9<br>0.08126 | sec per mg/kg/infusion cocaine<br>sec |
| | **H** | Post-reinforcement time-out responses vs. log cocaine dose | Linear regression | 27-38 | Slope<br>Y-intercept<br>$R^2$ | -17.6 to -7.4<br>5.2 to 11.0<br>0.1252 | time-out responses per mg/kg/infusion cocaine<br>time-out responses |
| | **I** | Earned reinforcements vs. log remifentanil dose | Nonlinear regression, second order polynomial | 15-34 | B0<br>B1<br>B2<br>$R^2$ | 17.34 to 21.38<br>-17.03 to -8.53<br>-7.71 to 3.05<br>0.1945 | mg/kg/infusion remifentanil<br>mg/kg/infusion remifentanil<br>mg/kg/infusion remifentanil |
| | **J** | Total active lever responses vs. log remifentanil dose | Nonlinear regression, second order polynomial | 15-34 | B0<br>B1<br>B2<br>$R^2$ | 82.77 to 106.8<br>-88.45 to -37.86<br>-37.22 to -9.45<br>0.1671 | mg/kg/infusion remifentanil<br>mg/kg/infusion remifentanil<br>mg/kg/infusion remifentanil |

| | | | | | | |
|---|---|---|---|---|---|---|
| **K** | Total inactive lever responses vs. log remifentanil dose | Linear regression | 15-34 | Slope<br><br>Y-intercept<br>$R^2$ | -13.15 to -6.58<br><br>7.41 to 15.24<br>0.1733 | lever response per mg/kg/infusion remifentanil<br>lever responses |
| **L** | Remifentanil consumed vs. log remifentanil dose | Nonlinear regression, exponential growth | 15-34 | Y0<br>k<br>Tau<br>Doubling Time<br>$R^2$ | 16.26 to 18.63<br>1.68 to 2.00<br>0.50 to 0.59<br>0.35 to 0.41<br>0.8791 | $(mg/kg/infusion\ remifentanil)^{-1}$<br>mg/kg/infusion remifentanil |
| **M** | Percent active lever responses vs. log remifentanil dose | Linear regression | 15-34 | Slope<br><br>Y-intercept<br>$R^2$ | 3.21 to 6.84<br><br>88.30 to 92.63<br>0.1506 | percent active lever responses per mg/kg/infusion remifentanil<br>percent active lever responses |
| **N** | Latency to first lever response vs. log remifentanil dose | Linear regression | 15-34 | Slope<br>Y-intercept<br>$R^2$ | -5.18 to 4.63<br>6.27 to 18.00<br>7.334e-005 | sec per mg/kg/infusion<br>sec |
| **O** | Latency to first earned reinforcement vs. log remifentanil dose | Linear regression | 15-34 | Slope<br>Y-intercept<br>$R^2$ | -48.00 to -15.53<br>16.79 to 55.41<br>0.08246 | sec per mg/kg/infusion<br>sec |
| **P** | Post-reinforcement time-out responses vs. log remifentanil dose | Linear regression | 15-34 | Slope<br>Y-intercept<br>$R^2$ | -14.50 to -5.29<br>12.86 to 23.85<br>0.09676 | responses per mg/kg/infusion<br>cue responses |

CI, confidence interval; sec, seconds.

#### Table S4. Curve parameters and statistical analysis of data presented in Figure 3 -- Supporting Figure 4.

| Figure | Panel | Experiment Description | F/t Statistics[1] | Multiple Comparisons[2] | N | Curve Fit | Parameter | 95% CI | Units |
|---|---|---|---|---|---|---|---|---|---|
| 3 | A | Active and inactive lever responses vs. extinction session number, **Cocaine** | $F_{Session}$(3.16, 156.6)=34.4, p<0.0001<br>$F_{Lever}$(1,64)=38.6, p<0.0001<br>$F_{Interaction}$(21, 1042)=15.5, p<0.0001<br><br>Geisser-Greenhouse's ε=0.15 | *p<0.001-p<0.05 Active vs. Inactive Lever | 14-32 | | | | |
| | B | Total lever presses vs. extinction session number, **Cocaine** | | | 14-32 | Nonlinear regression, exponential one-phase decay | Y0<br>Plateau<br>K<br>Half Life<br>Tau<br>$R^2$ | 202.1 to 296.7<br>36.19 to 43.80<br>0.4977 to 0.8295<br>0.8356 to 1.393<br>1.206 to 2.009<br>0.4218 | lever responses<br>lever responses<br>$(session)^{-1}$<br>session<br>session |
| | C, left | Lever discrimination: Active SA vs first extinction session, **Cocaine** | W = -458, p***<0.0001 | | 31 | | | | |
| | C, middle | Lever discrimination: First vs. Last extinction session, **Cocaine** | W = -367, p***<0.0001 | | 29 | | | | |
| | C, right | Lever discrimination: Last extinction session vs. Reinstatement session, **Cocaine** | W = 339, p***<0.0001 | | 29 | | | | |
| | D | Active and inactive lever responses by session type, **Cocaine** | $F_{Session}$(1.86,109.8)=76.0, p<0.0001<br>$F_{Lever}$(1,64)=98.5, p<0.0001<br>$F_{Interaction}$(2,118)=93.9, p<0.0001<br><br>Geisser-Greenhouse's ε=0.93 | ***p<0.0001, Active Lever SA vs. Active Lever Extinction; Active Lever Extinction vs. Active Lever Reinstatement<br><br>###p<0.0001 Active vs. Inactive Lever | 29-32 | | | | |
| | E | Active and inactive lever responses vs. extinction session number, **Remifentanil** | $F_{Session}$(3.78, 176.7)=60.9, p<0.0001<br>$F_{Lever}$(1,56)=37.2, p<0.0001<br>$F_{Interaction}$(20, 936)=30.5, p<0.0001<br><br>Geisser-Greenhouse's ε=0.19 | *p<0.001-p<0.05 Active vs. Inactive Lever | 22-28 | | | | |
| | F | Total lever presses vs. extinction session number, | | | 22-28 | Nonlinear regression, exponential | Y0<br>Plateau<br>K | 300.2 to 400.8<br>36.63 to 48.19<br>0.4086 to 0.6137 | lever responses<br>lever responses<br>$(session)^{-1}$ |

| | | | | | | | | |
|---|---|---|---|---|---|---|---|---|
| | **Remifentanil** | | | | one-phase decay | Half Life<br>Tau<br>$R^2$ | 1.129 to 1.696<br>1.630 to 2.447<br>0.4967 | session<br>session |
| **G, left** | Lever discrimination: Active SA vs first extinction session, **Remifentanil** | W = -340, p***<0.0001 | | 28 | | | | |
| **G, middle** | Lever discrimination: First vs. Last extinction session, **Remifentanil** | W = -153, p*<0.0115 | | 22 | | | | |
| **G, right** | Lever discrimination: Last extinction session vs. Reinstatement session, **Remifentanil** | W = 139, p*<0.0224 | | 22 | | | | |
| **H** | Active and inactive lever responses by session type, **Remifentanil** | $F_{Session}$(1.98,85.15)=20.4, p<0.0001<br>$F_{Lever}$(1,58)=67.2, p<0.0001<br>$F_{Interaction}$(2,868)=21.7, p<0.0001<br><br>Geisser-Greenhouse's ε=0.99 | ***p<0.0001, Active Lever SA vs. Active Lever Extinction; Active Lever Extinction vs. Active Lever Reinstatement<br><br>###p<0.0001 Active vs. Inactive Lever | 22-29 | | | | |

[1]F statistics from two-way, mixed effects analyses and W statistics from two-tailed, Wilcoxen Matched-Pairs Test; [2]*Post-hoc* Sidak tests for panels A and E, and Tukey's multiple comparisons tests for panels D and H. Sphericity was not assumed and the analyses were corrected using the Geisser-Greenhouse epsilon (ε) hat method; CI, confidence internal; SD, standard deviation; n.s., not significant.

#### Table S5. Correlation matrix for cocaine taking and cocaine seeking assessments – Supporting Figure 5.

| | | Acquisition Sessions Required | Reinforce-ments | Active Lever Responding | Inactive Lever Responding | Time-out Responding | Accuracy | Latency First Response | Latency First Reward | Early Extinction | Late Extinction | Reinstate -ment | Extinction Sessions Required |
|---|---|---|---|---|---|---|---|---|---|---|---|---|---|
| Pearson Correlation Coefficients | Acquisition Sessions Required | 1 | -0.037 | -0.073 | 0.242 | 0.218 | -0.237 | 0.337 | 0.427 | -0.135 | 0.223 | -0.211 | 0.316 |
| | Reinforcements | -0.037 | 1 | 0.915 | 0.23 | 0.359 | 0.312 | -0.38 | -0.579 | 0.636 | 0.101 | 0.32 | -0.127 |
| | Active Lever Responding | -0.073 | 0.915 | 1 | 0.143 | 0.514 | 0.373 | -0.41 | -0.59 | 0.619 | 0.038 | 0.316 | -0.181 |
| | Inactive Lever Responding | 0.242 | 0.23 | 0.143 | 1 | 0.322 | -0.768 | -0.223 | -0.016 | 0.11 | 0.245 | 0.055 | 0.232 |
| | Time-out Responding | 0.218 | 0.359 | 0.514 | 0.322 | 1 | -0.099 | -0.079 | -0.054 | 0.145 | 0.16 | 0.084 | 0.19 |
| | Accuracy | -0.237 | 0.312 | 0.373 | -0.768 | -0.099 | 1 | 0.01 | -0.291 | 0.223 | -0.182 | 0.028 | -0.298 |
| | Latency first response | 0.337 | -0.38 | -0.41 | -0.223 | -0.079 | 0.01 | 1 | 0.793 | -0.244 | -0.129 | -0.013 | -0.046 |
| | Latency first reward | 0.427 | -0.579 | -0.59 | -0.016 | -0.054 | -0.291 | 0.793 | 1 | -0.497 | -0.067 | -0.018 | 0.113 |
| | Early Extinction | -0.135 | 0.636 | 0.619 | 0.11 | 0.145 | 0.223 | -0.244 | -0.497 | 1 | -0.033 | 0.249 | -0.183 |
| | Late Extinction | 0.223 | 0.101 | 0.038 | 0.245 | 0.16 | -0.182 | -0.129 | -0.067 | -0.033 | 1 | -0.101 | 0.741 |
| | Reinstatement | -0.211 | 0.32 | 0.316 | 0.055 | 0.084 | 0.028 | -0.013 | -0.018 | 0.249 | -0.101 | 1 | -0.107 |
| | Extinction Sessions Required | 0.316 | -0.127 | -0.181 | 0.232 | 0.19 | -0.298 | -0.046 | 0.113 | -0.183 | 0.741 | -0.107 | 1 |
| Significance level (one-tailed) | Acquisition Sessions Required | | 0.414 | 0.333 | 0.075 | 0.098 | 0.079 | 0.021 | 0.004 | 0.212 | 0.093 | 0.105 | 0.029 |
| | Reinforcements | 0.414 | | 0 | 0.082 | 0.013 | 0.028 | 0.009 | 0 | 0 | 0.274 | 0.025 | 0.224 |
| | Active Lever Responding | 0.333 | 0 | | 0.195 | 0 | 0.011 | 0.005 | 0 | 0 | 0.411 | 0.027 | 0.138 |
| | Inactive Lever Responding | 0.075 | 0.082 | 0.195 | | 0.025 | 0 | 0.089 | 0.461 | 0.255 | 0.069 | 0.371 | 0.081 |
| | Time-out Responding | 0.098 | 0.013 | 0 | 0.025 | | 0.277 | 0.319 | 0.373 | 0.192 | 0.169 | 0.309 | 0.126 |
| | Accuracy | 0.079 | 0.028 | 0.011 | 0 | 0.277 | | 0.475 | 0.038 | 0.089 | 0.136 | 0.433 | 0.034 |
| | Latency first response | 0.021 | 0.009 | 0.005 | 0.089 | 0.319 | 0.475 | | 0 | 0.07 | 0.22 | 0.47 | 0.393 |
| | Latency first reward | 0.004 | 0 | 0 | 0.461 | 0.373 | 0.038 | 0 | | 0.001 | 0.345 | 0.457 | 0.249 |
| | Early Extinction | 0.212 | 0 | 0 | 0.255 | 0.192 | 0.089 | 0.07 | 0.001 | | 0.429 | 0.085 | 0.158 |
| | Late Extinction | 0.093 | 0.274 | 0.411 | 0.069 | 0.169 | 0.136 | 0.22 | 0.345 | 0.429 | | 0.291 | 0 |
| | Reinstatement | 0.105 | 0.025 | 0.027 | 0.371 | 0.309 | 0.433 | 0.47 | 0.457 | 0.085 | 0.291 | | 0.284 |
| | Extinction Sessions Required | 0.029 | 0.224 | 0.138 | 0.081 | 0.126 | 0.034 | 0.393 | 0.249 | 0.158 | 0 | 0.284 | |

#### Table S6. Correlation matrix for remifentanil taking and remifentanil seeking assessments – Supporting Figure 5.

| | | Acquisition Sessions Required | Reinforce-ments | Active Lever Responding | Inactive Lever Responding | Time-out Responding | Accuracy | Latency First Response | Latency First Reward | Early Extinction | Late Extinction | Reinstate-ment |
|---|---|---|---|---|---|---|---|---|---|---|---|---|
| Pearson Correlation Coefficients | Acquisition Sessions Required | 1 | 0.261 | 0.18 | -0.201 | -0.049 | 0.272 | 0.019 | -0.049 | -0.306 | -0.388 | -0.349 |
| | Reinforcements | 0.261 | 1 | 0.94 | 0.327 | 0.13 | 0.198 | -0.474 | -0.485 | 0.178 | 0.111 | -0.045 |
| | Active Lever Responding | 0.18 | 0.94 | 1 | 0.395 | 0.422 | 0.163 | -0.396 | -0.425 | 0.318 | 0.156 | -0.037 |
| | Inactive Lever Responding | -0.201 | 0.327 | 0.395 | 1 | 0.07 | -0.783 | -0.186 | -0.138 | 0.387 | 0.395 | 0.273 |
| | Time-out Responding | -0.049 | 0.13 | 0.422 | 0.07 | 1 | 0.15 | 0.101 | -0.005 | 0.268 | 0.088 | -0.036 |
| | Accuracy | 0.272 | 0.198 | 0.163 | -0.783 | 0.15 | 1 | 0.005 | -0.105 | -0.238 | -0.365 | -0.287 |
| | Latency first response | 0.019 | -0.474 | -0.396 | -0.186 | 0.101 | 0.005 | 1 | 0.853 | -0.188 | -0.143 | -0.114 |
| | Latency first reward | -0.049 | -0.485 | -0.425 | -0.138 | -0.005 | -0.105 | 0.853 | 1 | -0.153 | -0.064 | -0.124 |
| | Early Extinction | -0.306 | 0.178 | 0.318 | 0.387 | 0.268 | -0.238 | -0.188 | -0.153 | 1 | 0.722 | 0.36 |
| | Late Extinction | -0.388 | 0.111 | 0.156 | 0.395 | 0.088 | -0.365 | -0.143 | -0.064 | 0.722 | 1 | 0.601 |
| | Reinstatement | -0.349 | -0.045 | -0.037 | 0.273 | -0.036 | -0.287 | -0.114 | -0.124 | 0.36 | 0.601 | 1 |
| Significance level (one-tailed) | Acquisition Sessions Required | | 0.071 | 0.158 | 0.131 | 0.394 | 0.063 | 0.458 | 0.394 | 0.042 | 0.013 | 0.023 |
| | Reinforcements | 0.071 | | 0 | 0.03 | 0.233 | 0.131 | 0.002 | 0.002 | 0.156 | 0.266 | 0.399 |
| | Active Lever Responding | 0.158 | 0 | | 0.01 | 0.006 | 0.178 | 0.01 | 0.006 | 0.033 | 0.189 | 0.418 |
| | Inactive Lever Responding | 0.131 | 0.03 | 0.01 | | 0.348 | 0 | 0.146 | 0.218 | 0.012 | 0.01 | 0.059 |
| | Time-out Responding | 0.394 | 0.233 | 0.006 | 0.348 | | 0.198 | 0.285 | 0.49 | 0.063 | 0.309 | 0.419 |
| | Accuracy | 0.063 | 0.131 | 0.178 | 0 | 0.198 | | 0.49 | 0.278 | 0.088 | 0.017 | 0.05 |
| | Latency first response | 0.458 | 0.002 | 0.01 | 0.146 | 0.285 | 0.49 | | 0 | 0.144 | 0.209 | 0.26 |
| | Latency first reward | 0.394 | 0.002 | 0.006 | 0.218 | 0.49 | 0.278 | 0 | | 0.193 | 0.359 | 0.242 |
| | Early Extinction | 0.042 | 0.156 | 0.033 | 0.012 | 0.063 | 0.088 | 0.144 | 0.193 | | 0 | 0.028 |
| | Late Extinction | 0.013 | 0.266 | 0.189 | 0.01 | 0.309 | 0.017 | 0.209 | 0.359 | 0 | | 0.001 |
| | Reinstatement | 0.023 | 0.399 | 0.418 | 0.059 | 0.419 | 0.05 | 0.26 | 0.242 | 0.028 | 0.001 | |

**Table S7. Factor loadings of variables in exploratory factor analyses – Supporting Figure 6.**

| | Within-Drug Cocaine | | Within-Drug Remifentanil | | Global | |
|---|---|---|---|---|---|---|
| **Variable** | **Factor 1** | **Factor 2** | **Factor 1** | **Factor 2** | **Factor 1** | **Factor 2** |
| Acquisition sessions required | -0.072 | 0.270 | 0.179 | -0.296 | -0.326 | 0.147 |
| Reinforcements* | 0.952 | 0.011 | 0.941 | -0.054 | 0.925 | 0.075 |
| Active lever responding* | 0.964 | -0.073 | 0.999 | -0.004 | 0.991 | 0.060 |
| Inactive lever responding* | 0.224 | 0.945 | 0.399 | 0.889 | 0.321 | 0.885 |
| Percent time-out responses* | 0.454 | 0.249 | 0.419 | -0.091 | 0.551 | 0.121 |
| Lever accuracy* | 0.323 | -0.889 | 0.159 | -0.951 | 0.522 | -0.818 |
| Latency to first lever response* | -0.433 | -0.136 | -0.397 | -0.052 | -0.417 | -0.145 |
| Latency to first reinforcement* | -0.613 | 0.122 | -0.426 | 0.033 | -0.660 | 0.068 |
| Early extinction lever responding | 0.649 | -0.035 | 0.319 | 0.314 | 0.527 | 0.204 |
| Late extinction lever responding | 0.072 | 0.253 | 0.158 | 0.408 | 0.229 | 0.344 |
| Reinstatement lever responding | 0.311 | 0.010 | -0.035 | 0.325 | 0.233 | 0.191 |
| Extinction sessions required | -0.150 | 0.296 | N/A | N/A | N/A | N/A |
| **Eigenvalue**[1] | 3.728 | 2.562 | 3.396 | 2.673 | 4.195 | 2.004 |
| **% Variance**[1] | 31.069 | 21.351 | 30.871 | 24.300 | 38.140 | 18.219 |
| **Cumulative %**[1] | 52.420 | | 55.171 | | 56.359 | |
| **N** | 38 | | 34 | | 72 | |

*Averaged values from the 0.1, 0.3, 0.5, and 1.0 mg/kg/infusion sessions for mice in the cocaine paradigm and from the 0.01, 0.03, 0.1, 0.3, and 1.0 mg/kg/infusion sessions for mice in the remifentanil paradigm.[1]Based on Initial eigenvalues.

**Table S8. Component loadings of variables in novelty-induced open field measures principal component analysis – Supporting Figure 6.**

| Variable | Factor 1 | Factor 2 | Factor 3 | Factor 4 | Factor 5 |
|---|---|---|---|---|---|
| Distance | -0.474 | -0.268 | 0.018 | -0.414 | 0.473 |
| Vertical Episodes | -0.421 | -0.351 | 0.131 | 0.822 | 0.004 |
| Center Time | -0.222 | 0.613 | 0.737 | 0.033 | 0.171 |
| Rest Time | 0.473 | -0.234 | 0.137 | 0.142 | 0.780 |
| Stereotypic Episodes | -0.302 | 0.543 | -0.645 | 0.192 | 0.373 |
| Velocity | -0.484 | -0.280 | 0.067 | -0.307 | -0.015 |
| **Eigenvalues**[1] | 3.74 | 1.01 | 0.83 | 0.31 | 0.16 |
| **% Variance**[1] | 61.44 | 16.50 | 13.54 | 5.16 | 2.66 |
| **Cumulative %**[1] | | | 99.3 | | |
| **N** | | | 65 | | |

[1]Based on Initial eigenvalues.

**Table S9. Component loadings of variables in drug taking principal component analyses – Supporting Figure 6.**

| Variable | Within-Drug Cocaine | | | | | Within-Drug Remifentanil | | | | |
|---|---|---|---|---|---|---|---|---|---|---|
| | **Factor 1** | **Factor 2** | **Factor 3** | **Factor 4** | **Factor 5** | **Factor 1** | **Factor 2** | **Factor 3** | **Factor 4** | **Factor 5** |
| Acquisition sessions required | 0.144 | 0.420 | 0.400 | -0.593 | 0.528 | -0.114 | -0.374 | 0.122 | -0.736 | 0.534 |
| Reinforcements | -0.471 | 0.157 | 0.243 | -0.302 | -0.448 | -0.519 | -0.082 | 0.152 | -0.196 | -0.424 |
| Active lever responding | -0.504 | 0.136 | 0.254 | 0.049 | -0.231 | -0.514 | -0.056 | 0.343 | -0.013 | -0.247 |
| Inactive lever responding | -0.063 | 0.585 | -0.365 | -0.115 | -0.260 | -0.231 | 0.609 | 0.234 | -0.164 | 0.020 |
| Percent time-out responses | -0.245 | 0.414 | 0.259 | 0.708 | 0.368 | -0.135 | -0.116 | 0.600 | 0.542 | 0.508 |
| Lever accuracy | -0.194 | -0.473 | 0.49 | -0.052 | 0.033 | -0.051 | -0.679 | -0.046 | 0.201 | -0.276 |
| Latency to first lever response | 0.428 | 0.108 | 0.444 | 0.130 | -0.307 | 0.429 | -0.067 | 0.489 | -0.141 | -0.213 |
| Latency to first reinforcement | 0.467 | 0.176 | 0.279 | 0.146 | -0.410 | 0.442 | 0.012 | 0.435 | -0.195 | -0.307 |
| **Eigenvalues[1]** | 3.28 | 2.17 | 1.59 | 0.59 | 0.43 | 3.08 | 1.98 | 1.41 | 0.96 | 0.60 |
| **% Variance[1]** | 39.66 | 26.22 | 19.23 | 7.15 | 5.14 | 37.41 | 24.00 | 17.15 | 11.65 | 7.31 |
| **Cumulative %[1]** | | | 97.4 | | | | | 97.5 | | |
| **N** | | | 31 | | | | | 34 | | |

[1]Based on Initial eigenvalues.

**Table S10. Linear regression models for drug seeking – Supporting Figure 6.**

| | | Within-Drug Cocaine | | | Within-Drug Remifentanil | | |
|---|---|---|---|---|---|---|---|
| | | Early Extinction | Late Extinction | Reinstatement | Early Extinction | Late Extinction | Reinstatement |
| Linear Regression Coefficients | Intercept | 543.1796*** | 488.63*** | 50.689*** | 490.77*** | 774.511*** | 65.064*** |
| | Drug-taking PC 1 | -75.1991*** | N/I | N/I | -52.70** | N/I | N/I |
| | Drug-taking PC 2 | 29.2839 | N/I | N/I | N/I | 139.643** | 8.610* |
| | Drug-taking PC 3 | 21.0105 | N/I | 2.394 | 79.06** | N/I | N/I |
| | Drug-taking PC 4 | -66.7308* | N/I | N/I | N/I | N/I | N/I |
| | Drug-taking PC 5 | N/I | N/I | -29.606** | N/I | N/I | N/I |
| | Open field PC 1 | N/I | N/I | N/I | N/I | N/I | N/I |
| | Open field PC 2 | N/I | N/I | N/I | 100.53* | N/I | N/I |
| | Open field PC 3 | -120.2241** | 113.89 | N/I | N/I | N/I | N/I |
| | Open field PC 4 | 54.6585 | N/I | N/I | 70.34 | N/I | N/I |
| | Open field PC 5 | 162.0896** | N/I | N/I | 199.17** | 392.030* | 43.27** |
| | Cocaine-induced hyperlocomotion¥ | -0.7613* | N/I | N/I | N/A | N/A | N/A |
| | Remifentanil-induced hyperlocomotion¥ | N/A | N/A | N/A | N/I | -8.848. | N/I |
| Performance | **Residual standard error** | 91.19 | 218.8 | 24.43 | 136.1 | 349.8 | 28.06 |
| | **Multiple R-squared** | 0.7697 | 0.1073 | 0.369 | 0.6829 | 0.4321 | 0.4293 |
| | **Adjusted R-squared** | 0.6469 | 0.06677 | 0.309 | 0.6074 | 0.358 | 0.3817 |
| | **F statistic** | 6.266 | 2.646 | 6.142 | 9.044 | 5.833 | 9.025 |
| | **Degrees of Freedom** | 15 | 22 | 21 | 21 | 23 | 24 |
| | **P value** | 0.00118 | 0.118 | 0.00794 | 0.000105 | 0.00407 | 0.00120 |

Significance codes: *$p<0.05$, **$p<0.01$, ***$p<0.001$
Abbreviations: N/I, not included in the model based on Akaike information criterion; N/A, not applicable.
[¥]Total distances value for 20 mg/kg (i.p.) cocaine and 1 mg/kg (i.p.) remifentanil.
